# P23H rhodopsin aggregation in the ER causes synaptic protein imbalance in rod photoreceptors

**DOI:** 10.1101/2024.10.18.619115

**Authors:** Samantha L. Thompson, Sophie M. Crowder, Maryam Hekmatara, Emily R. Sechrest, Wen-Tao Deng, Michael A. Robichaux

## Abstract

Rod photoreceptor neurons in the retina detect scotopic light through the visual pigment rhodopsin (Rho) in their outer segments (OS). Efficient Rho trafficking to the OS through the inner rod compartments is critical for long-term rod health. Given the importance of protein trafficking to the OS, less is known about the trafficking of rod synaptic proteins. Furthermore, the subcellular impact of Rho mislocalization on rod synapses (i.e., “spherules”) has not been investigated. In this study we used super-resolution and electron microscopies, along with proteomics, to perform a subcellular analysis of Rho synaptic mislocalization in P23H-Rho-RFP mutant mice. We discovered that mutant P23H-Rho-RFP protein mislocalized in distinct ER aggregations within the spherule cytoplasm, which we confirmed with AAV overexpression. Additionally, we found synaptic protein abundance differences in P23H-Rho-RFP mice. By comparison, Rho mislocalized along the spherule plasma membrane in WT and rd10 mutant rods, in which there was no synaptic protein disruption. Throughout the study, we also identified a network of ER membranes within WT rod presynaptic spherules. Together, our findings indicate that photoreceptor synaptic proteins are sensitive to ER dysregulation.

**Summary Statement:** This study examines the impact of rhodopsin mislocalization on rod photoreceptor synaptic structures and synaptic protein levels using P23H rhodopsin and other retinitis pigmentosa mouse models.

## Introduction

In the retina, rod photoreceptor neurons detect dim light through the photoactivation of the rod-specific G-protein coupled receptor rhodopsin (Rho). Rho and other visual proteins are densely loaded into stacked membrane discs in the rod outer segment (OS) compartment, which is joined to the inner segment (IS) by a narrow connecting cilium (CC). Mammalian rods have compartmentalized cell bodies preceding the presynaptic terminals (spherules), which form synapses with downstream retinal neurons (Townes-Anderson et al., 1988). Proper Rho protein trafficking to the OS is absolutely essential for long-term rod stability and retinal health (Sung et al., 1994; Lem et al., 1999). Since new OS membrane discs are continuously formed in rods, Rho protein, which is synthesized throughout the cell body and IS, must be constantly fluxed unidirectionally into the OS through various coordinated trafficking mechanisms. Any disruption to the unidirectional flow of Rho into the OS causes Rho mislocalization, which is the typical subcellular outcome of blinding rod diseases caused either by a genetic mutation (Guo et al., 2022; Hagstrom et al., 1999) or retinal detachment (Fariss et al., 1997; Fisher et al., 2005). Despite the central role of OS protein trafficking in rods, a cellular trafficking system in rods is required to supply and maintain their presynaptic spherules; however, little is known about these trafficking mechanisms and how they might be affected by Rho mislocalization.

Rod presynaptic spherules are located in the outer plexiform layer (OPL) of the retina and contain a tetrad of postsynaptic invaginating neurites (Behrens et al., 2016; Torten et al., 2023). Each rod spherule features a single synaptic ribbon, an electron dense structure that organizes synaptic vesicles for glutamate release in the dark (Moser et al., 2020). Most mouse rods have stereotypical R1 spherules that are connected to the cell body through an axon (or “internal fiber”, (Carter-Dawson and Lavail, 1979)), while fewer rods have R2 spherules that are contiguous with the cell bodies (Fig. 1 A; (Li et al., 2016)). Critically, rod spherules contain essential proteins that are either structural elements of the synaptic ribbon, like BASSOON, or proteins that localize at the synaptic cleft to form stabilizing trans-synaptic protein complexes, including ELFN1, Dystrophin and Dystroglycan (Furukawa et al., 2020). Disruptions to these rod synaptic proteins lead to functional and structural defects in spherules and often cause irreparable synaptic miswiring (Dick et al., 2003; Omori et al., 2012).

**Figure 1.**
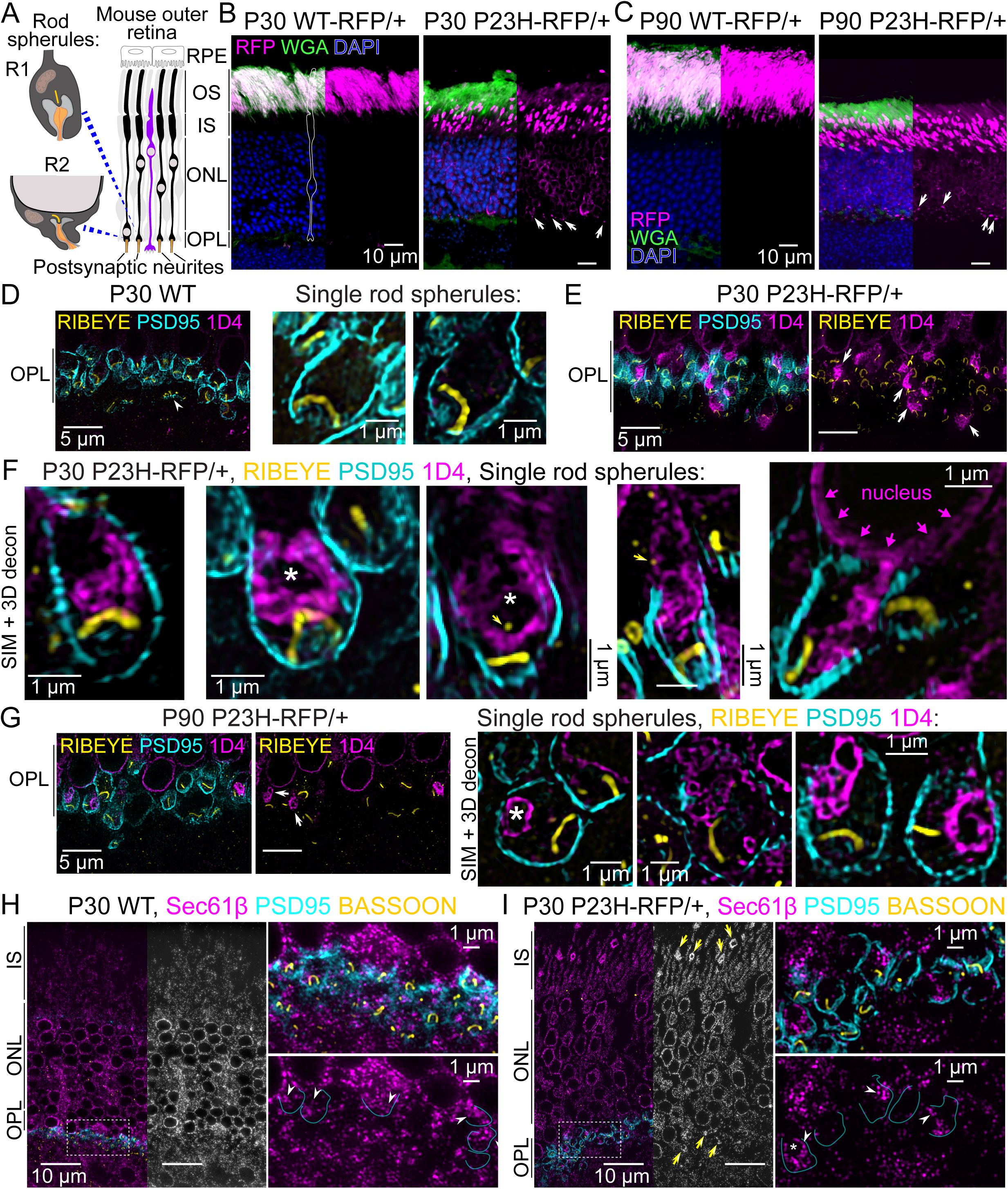
Mutant P23H-hRho-RFP protein is mislocalized within the cytoplasm of rod photoreceptor presynaptic spherules. (A) Diagram depicting the layers of the mouse outer retina (RPE = retinal pigment epithelium, OS = outer segment, IS = inner segment, ONL = outer nuclear layer, OPL = outer plexiform layer) and the two types of rod spherules (R1, top; R2, bottom). Rod photoreceptors are black, and the cone photoreceptor is purple. Spherule illustrations were based on (Li et al., 2016). (B, C) Confocal z-projections of *WT-RFP/+* and *P23H-RFP/+* retinal cryosections at age (B) P30 and (C) P90. RFP fluorescence is magenta, and sections were co-stained with WGA to label OS membranes (green) and DAPI to label nuclei (blue). White arrows = mislocalized RFP in the *P23H-RFP/+* OPLs. (D) SIM super-resolution z-projections with 3D deconvolution of the OPL from a P30 WT retina. In the images, RIBEYE (yellow) and PSD95 (cyan) immunolabeled rod spherules are aligned in the OPL above a cone pedicle (white arrowhead). No 1D4 Rho labeling (magenta) is present in the WT OPL. In single spherule examples, which are all sub-stack SIM z-projections, the RIBEYE+ ribbons are horseshoe-shaped structures in the lower region of the spherules. (E, F) SIM images of the OPL in a *P23H-RFP/+* retina at age P30 with the same immunolabeling as in (D). In (E) the left images include all labeling, and the right shows the 1D4 and RIBEYE channels without PSD95. Accumulations of 1D4 immunolabeling in the OPL were localized near the synaptic ribbons (white arrows). (F) Single spherule SIM examples in *P23H-RFP/+* P30 retinas. 1D4+ Rho accumulations are localized in the cytoplasm of the spherules, typically above the ribbon. In the second SIM + 3D decon. image, a gap in the aggregated 1D4+ fluorescence is indicated with a white asterisk. The far-right example is a R2-type mutant spherule with 1D4 fluorescence that surrounds the nucleus (magenta arrows) and extends into the spherule cytoplasm. (G) SIM images of *P23H-RFP/+* retinas at age P90 with the same immunolabeling as (D-F). Images of the OPL with and without PSD95 show 1D4+ OPL accumulations (white arrows) similar to those at P30. In single spherule examples, cytoplasmic 1D4+ aggregates surround gaps in fluorescence (white asterisk). (H, I) SIM z-projection images of (H) P30 WT and (I) P30 *P23H- RFP/+* retinas immunolabeled for Sec61β (ER-marker, magenta), PSD95 (cyan), and BASSOON (yellow). Sec61β is localized throughout the layers of the photoreceptors (left), and in magnified images, the PSD95 rod spherule border is annotated in select rod spherules to demonstrate Sec61β+ ER fluorescence within individual WT and *P23H-RFP/+* rod spherules (white arrowheads). In the *P23H-RFP/+* image in (G), ER aggregations are labeled with yellow arrows. Throughout the figure scale bars match adjacent images when not labeled.

Therefore, rods must utilize a secretory system to maintain their presynaptic spherules that operates in concert with constant protein delivery to the OS. In mouse models retinitis pigmentosa (RP) of photoreceptor degeneration, including for the well-characterized misfolding P23H-Rho mutation (Kaushal and Khorana, 1994; Saliba et al., 2002; Sung et al., 1991), Rho is mislocalized not only to the rod IS and cell body but also the OPL (Barhoum et al., 2008; Hagstrom et al., 1999; Roof et al., 1994). In the *P23H-hRho-Tag-RFP-T* mouse, mutant P23H- hRho-RFP protein mislocalized and accumulated throughout the ER in the IS, ONL, but also in the OPL (Robichaux et al., 2022). Rho mislocalization in the OPL has also been demonstrated in RP dog and human retinas (Beltran et al., 2006; Fariss et al., 2000; Milam et al., 1998), as well as after retinal detachment (Fariss et al., 1997; Fisher et al., 2005); however, the impact of Rho OPL mislocalization on rod spherule structure and synaptic protein trafficking and turnover has never been investigated.

Here, we performed a detailed subcellular analysis of Rho mislocalization in rod spherules. Using *P23H-RFP* mice, we found that P23H-hRho-RFP mutant protein accumulated within the ER of the spherule cytoplasm, as opposed to membrane mislocalization observed in rd10 RP mutant rods or in overloaded WT rods. Mutant P23H-Rho ER aggregation in *P23H- RFP/+* rods interfered with normal rod synaptic protein levels. Combined with evidence of ER membranes in WT rod spherules, our findings indicate that an ER-based secretory system may regulate presynaptic protein content within mouse rod spherules.

## Results

### Mutant P23H-Rho-RFP protein is mislocalized within the cytoplasm of rod presynaptic spherules

To exclude the possibility that Rho mislocalization in *P23H-hRho-TagRFP-T* (hereafter *P23H-RFP*) heterozygous knockin mouse retinas is the outcome of the P23H-Rho misfolding mutation rather than an effect of the C-terminal TagRFP-T fusion tag, a new *WT-hRho-TagRFP-T* mouse line (hereafter “*WT-RFP*” in reference to the knockin mouse) was generated with a restored WT P23 residue in the *hRho-RFP-TagRFP-T* knockin allele. As in the *P23H-RFP* mice, an additional 1D4 sequence is included at the C-terminus of the *WT-RFP* allele. In *WT-RFP/+* heterozygous retinas, WT-hRho-TagRFP-T fusion protein (hereafter WT-hRho-RFP) was localized exclusively in the OS layer (Fig. 1B,C). By comparison, in *P23H-RFP/+* heterozygous mouse retinas, P23H-hRho-TagRFP-T protein (abbreviated P23H-hRho-RFP) was mislocalized throughout the IS, ONL, and OPL at ages P30 and P90 (Fig. 1B,C) as described previously (Robichaux et al., 2022). The normal OS localization of WT-hRho-RFP demonstrates that Rho mislocalization in *P23H-RFP/+* rods is triggered by the P23H-Rho mutation, not the TagRFP-T fusion tag. Additionally, western blotting with the 1D4 Rho monoclonal antibody (Molday and MacKenzie, 1983) was used to demonstrate a ∼65 kDa band corresponding to the WT-hRho-RFP protein in *WT-RFP/+* retinas (Fig. S1A, magenta arrow). A deglycosylation assay showed a lower molecular weight band for Rho which confirmed that WT-hRho-RFP was glycosylated just like endogenous mouse WT Rho protein and WT Rho-GFP-1D4 protein from the *Rho-GFP- 1D4/+* mouse (Haggerty et al., 2024; Robichaux et al., 2022) (Fig. S1B, magenta arrow). ONL density was quantified to compare P30 WT retinas to P30 *WT-RFP/+* retinas, revealing no significant difference (Fig. S1C), which demonstrates that *WT-RFP/+* rods are stable at this age. However, there was ONL photoreceptor nuclei loss in P180 *WT-RFP/+* retinas indicating late-stage instability. In a separate analysis, ONL nuclei counts in *WT-RFP/+* retinas were significantly higher than those in age-matched *P23H-RFP/+* retinas at both P30 and P90 (Fig. S1D).

Next, SIM super resolution microscopy was used to closely examine the mislocalization of P23H-hRho-RFP mutant protein in the OPL of *P23H-RFP/+* retinas. At P30, *P23H-RFP/+* retinas were shown to be in an early stage of degeneration, before significant photoreceptor loss occurred (Robichaux et al., 2022). In the previous study, we focused on characterizing the mislocalized OPL Rho through co-labeling with ER markers. In the current study, we investigated P23H-Rho-RFP mislocalization within OPL using rod presynaptic spherule markers. *P23H-RFP/+* and age-matched WT retinas were co-immunolabeled with the 1D4 monoclonal Rho antibody, a RIBEYE antibody to visualize synaptic ribbons, and a PSD95 fluorescent nanobody to visualize the presynaptic spherule plasma membranes (Koulen et al., 1998). The samples were then imaged with SIM. In the OPL of P30 WT retinas, PSD95+ rod spherule plasma membranes encased single RIBEYE+ synaptic ribbons that typically appear as horseshoe-shaped structures (Fig. 1D, Fig. S1E). These rod spherules were distinguishable from cone presynaptic pedicles, which have multiple short ribbons. In P30 *P23H-RFP/+* retinas, bright aggregates of mislocalized Rho protein were located inside some of the rod spherules in the OPL (Fig. 1E, white arrows). Rho aggregations were detected in 28.3% ± 8.6% (mean ± SD, N=3 mice) of all P30 *P23H-RFP/+* rod spherules imaged with SIM. Upon closer observation of individual *P23H-RFP/+* rod spherules, mutant Rho aggregates were localized within the cytoplasm typically near the synaptic ribbons (Fig. 1F, Fig. S1F,G). In some spherules, there was a partial overlap in 1D4 and RIBEYE signals, and in some cases, the cytoplasmic P23H- hRho-RFP aggregates formed a swirling pattern of bright fluorescence surrounding a dark patch (white asterisks in Fig. 1F). RIBEYE+ synaptic ribbons appeared intact in most of the mutant *P23H-RFP/+* spherules examined. We also observed many cases of a small RIBEYE+ puncta in the cytoplasmic space in both WT and *P23H-RFP/+* spherules; in *P23H-RFP/+* spherules, these RIBEYE+ puncta often colocalized with the mutant P23H-hRho-RFP aggregates (Fig. 1F, Fig. S1G). These small RIBEYE+ puncta could represent recycling synaptic ribbons (Adly et al., 1999; Schmitz et al., 2012). While most 1D4+ P23H-hRho-RFP aggregates were observed as distinct puncta localized near a synaptic ribbon, in some cases, such as in R2 spherules, the 1D4+ P23H-hRho-RFP in the spherule cytoplasm formed a continuous network with the cytoplasm surrounding the adjacent cell body nucleus (Fig. 1F, magenta arrows).

The same SIM analysis was performed in *P23H-RFP/+* mice at age P90. At this age, *P23H-RFP/+* mice have significant photoreceptor degeneration, but the surviving rods were shown to be adaptive to the mutation, as the rate of degeneration and cell loss reduces after P90 (Robichaux et al., 2022). Here, in P90 mutant *P23H-RFP/+* rods, the mislocalized P23H- hRho-RFP protein accumulated in the OPL in a similar pattern as age P30 mutant rods (Fig. 1 G, Fig. S1 H,I) while there was no Rho staining in the OPL of WT P90 rods (Fig. S1J). Prominent Rho aggregations were detected in 24.4% ± 5.9% (mean ± SD, N=3 mice) of all P90 P23H-RFP/+ rod spherules imaged with SIM. In individual P90 mutant spherules, large P23H- hRho-RFP aggregations were again localized inside of the spherule cytoplasm, typically close to the ribbons, and were often observed in a swirling pattern around an empty gap in fluorescence. Thus, based on our SIM analysis, large cytoplasmic aggregations of mutant P23H-hRho-RFP protein occur in ∼25% of the spherules in the *P23H-RFP/+* rods at P30, and this phenotype persists in the surviving mutant rods at age P90.

Our SIM results, combined with our previous findings, suggest that the ER may be localized not only in mouse rod photoreceptor cell bodies but also in the presynaptic spherules. While previous studies localized the ER throughout the IS, ONL, and OPL in mouse retinas (Agrawal et al., 2017; Babai et al., 2010; Chen et al., 2015; Krizaj, 2005), we validated ER extension into the rod spherule cytoplasm using immunolabeling with Sec61β, an ER marker, and SIM. In both P30 WT and P23H-RFP/+ retinas, Sec61β ER labeling was distributed throughout all the photoreceptor inner subcompartments (Fig. 1H), but in the *P23H-RFP/+* retinas, larger Sec61β+ puncta were localized in the IS and OPL, corresponding to the fluorescent ER aggregates in those mutant rods. In higher magnification views of the OPL, ER immunofluorescence was clearly localized within the PSD95+ rod spherules in both genotypes (Fig. 1H,I), demonstrating that the ER network in rod photoreceptors extends from the IS to the rod presynaptic cytoplasm.

### P23H-Rho synaptic mislocalization does not cause ultrastructural defects in rod synaptic ribbons

Based on the observation throughout our SIM analysis that mislocalized P23H-Rho-RFP occupied a significant amount of the rod spherule cytoplasm, we hypothesized that these mislocalized cytoplasmic aggregates disrupted the rod presynaptic machinery on an ultrastructural level. In transmission electron microscopy (TEM) images of P30 WT mouse retinas, rod spherules have a distinct plasma membrane that surrounds the invaginating post-synaptic neurites, an electron dense presynaptic ribbon that extends from the active zone, a cytoplasm filled with synaptic vesicles, and a large mitochondrion (Fig. 2A). In many spherules, clusters of electron dense endocytic vesicles that are denser than other cytoplasmic synaptic vesicles were observed (Fig. 2A and Fig S2A, green arrowheads) (Fuchs et al., 2014). Additionally, more irregular membranes were found surrounding the spherule’s mitochondrion (Fig. 2A and Fig S2A, orange arrowheads), in a manner similar to ER membranes previously observed around mitochondria in other neurons (Wu et al., 2017) and in cat cone presynaptic pedicles (Lovas, 1971). Most P30 *P23H-RFP/+* mutant rod spherules had normal TEM ultrastructure, except in cases where electron dense bundles of folded membranes were observed within the spherule cytoplasm (Fig. 2B,C, orange arrows). The ultrastructure of these stacked membranes matches the semi-organized membrane stacks observed with TEM in the IS layer of *P23H-RFP/+* retinas (Robichaux et al., 2022), and they also match the localization and relative size of the P23H-Rho-RFP puncta observed in Figure 1. Therefore, we conclude that these are expanded stacks of ER membranes within *P23H-RFP/+* spherules that are filled with mislocalized and accumulated P23H-Rho-RFP protein.

**Figure 2.**
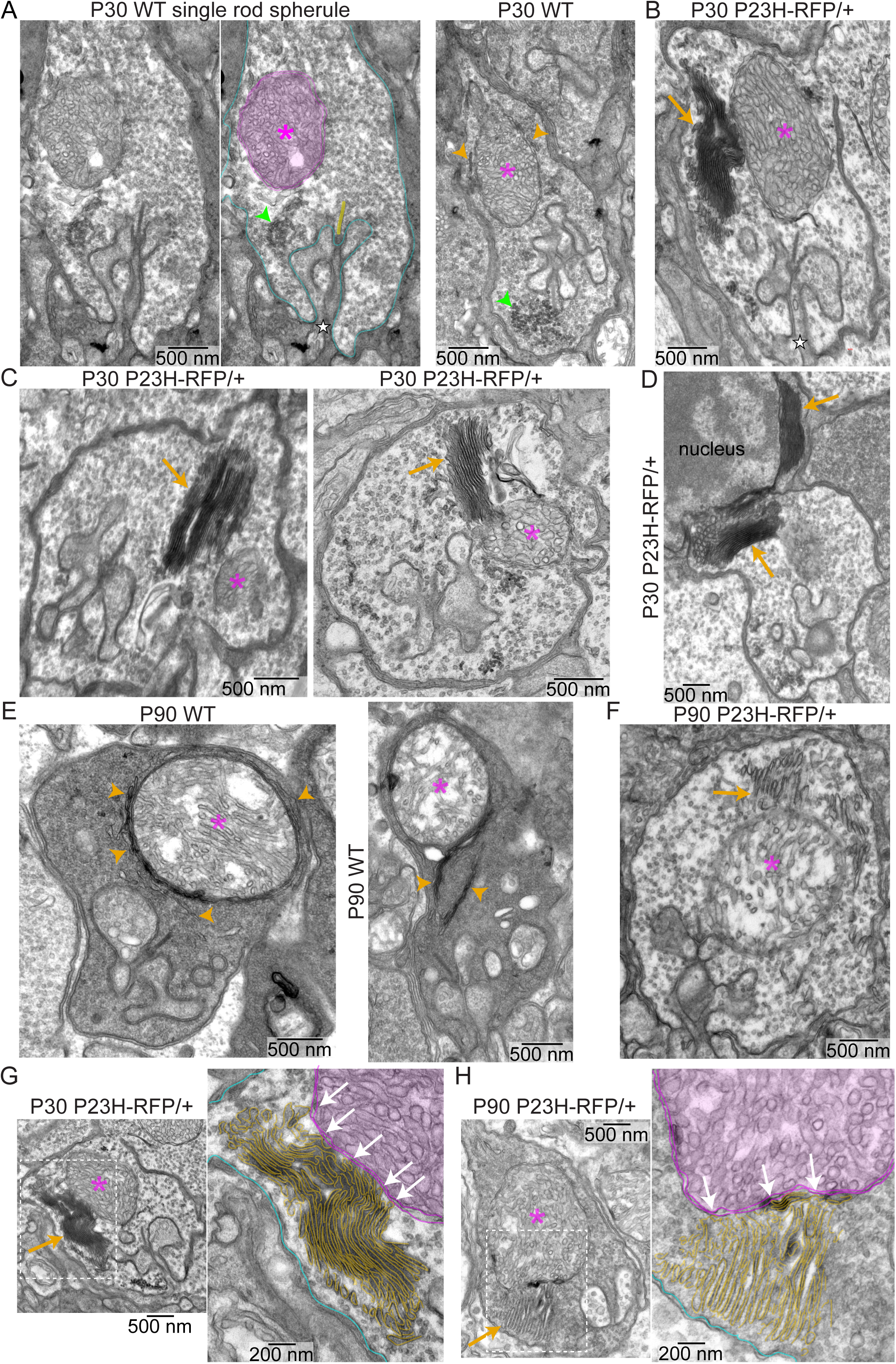
P23H-Rho-RFP mislocalization does not cause ultrastructural defects in rod synaptic ribbons. (A) TEM images of WT rod spherules at P30. Middle image is the annotated version of the left image (magenta asterisk = mitochondrion, cyan = spherule plasma membrane, yellow = synaptic ribbon, green arrowhead = endocytosed vesicles). In the right image, ER-like membranes (orange arrows) are wrapped around the mitochondrion. (B) TEM image of a *P23H-RFP/+* rod spherule at P30. ER-like membrane stacks (orange arrow) are localized near the mitochondrion (indicated with a magenta asterisk in all panels) and extend into the cytoplasm. (C) Additional TEM images of P30 *P23H-RFP/+* rod spherules with ER-like membrane stacks (orange arrow) associated with the spherule mitochondria. (D) TEM image of a P30 *P23H-RFP/+* R2 rod spherule with ER membrane stacks (orange arrow) from the cell body cytoplasm extending into the cytoplasm of the spherule. (E) TEM images of P90 WT spherules with ER membranes (orange arrows) surrounding the mitochondria and extending into the spherule cytoplasm. (F) TEM image of a P90 *P23H-RFP/+* spherule with expanded ER (orange arrow). (G, H) Magnified examples of ER-mitochondria contact sites (white arrows) in *P23H-RFP/+* spherules at ages (G) P30 and (H) P90. ER membranes are traced in gold, the mitochondrial membranes are traced in magenta, and the plasma membranes are traced in cyan.

In some *P23H-RFP/+* rod spherules, the dense ER membrane stacks were less compact or potentially discontinuous (Fig. S2B); however, in most spherules, some portion of the ER membrane stacks closely localized near the spherule’s mitochondrion. In all cases, the long continuous ER membranes were distinct from the electron dense endocytosed vesicles. In one P30 *P23H-RFP/+* R2 spherule, in which the spherule cytoplasm is continuous with the rod cell body, accumulated stacks of ER appeared to spill over from the cell body cytoplasm into the spherule cytoplasm (Fig. 2D).

Despite these large ER accumulations, there appeared to be no other major ultrastructural changes to the spherules or the synaptic ribbons in our P30 *P23H-RFP/+* TEM data. To confirm that the ribbon ultrastructure was unaffected, we measured the ribbon height and quantified the number of synaptic vesicles associated with the ribbons in TEM images of P30 WT and *P23H-RFP/+* spherules, and no significant differences were found. The average synaptic ribbon height in WT spherules was 349.4 nm ± 58.1 nm (average ± SD, N=53 ribbons, N=3 mice), while the average *P23H-RFP/+* ribbon height was 348.2 nm ± 85.1 nm (N=68 ribbons, N=3 mice). For WT spherules, the average number of ribbon-associated synaptic vesicles was 7.7 ± 1.8 per ribbon (N=46 ribbons, N=3 mice), while in *P23H-RFP/+* spherules, the average number of ribbon-associated synaptic vesicles was 7.4 SVs ± 2.4 SVs per ribbon (N=65 ribbons, N=3 mice).

The TEM ultrastructure of P90 WT rod spherules was similar to P30; however, there were more noticeable ER-like membranes surrounding the WT spherule mitochondria at this age (Fig. 2E). Based on our collective observations, we conclude that the irregular membranes surrounding the spherule mitochondria in P30 and P90 rod spherules are part of an endogenous network of ER within the spherule cytoplasm. In one P30 WT spherule TEM image, a portion of the axon was captured in a cross-section with the spherule from the same rod, and irregular membranes were observed within the axon shaft and possibly within a continuous network with the ER surrounding the spherule mitochondrion (Fig. S2C). In another image of a P90 WT R2 spherule, elongated ER-like membranes appeared to extend from the ER surrounding the nucleus into the spherule cytoplasm (Fig. S2D). P90 *P23H-RFP/+* mutant spherules also had expanded ER membranes that appeared tethered to the mitochondria (Fig. 2F). In higher-magnification TEM images of P30 and P90 *P23H-RFP/+* mutant spherules, the expanded ER membrane folds were traced and compared with mitochondrial membranes, and multiple possible ER-mitochondrion contact sites were found in each example (Fig. 2G,H).

### AAV overexpression of P23H-Rho and R135L-Rho in WT rods leads to synaptic mislocalization

To test if the P23H-Rho mutation causes mislocalization in the OPL of rod photoreceptors with a non-disease, healthy genetic background, AAVs expressing P23H-hRho-TagRFP-T (P23H-hRho-RFP) or R135L-hRho-EGFP (R135L-hRho-GFP) mutant Rho fusions driven by the rod-specific minimal mouse opsin promoter (Pawlyk et al., 2005) were subretinally injected in adult WT mice. The R135L-Rho mutant was analyzed alongside P23H-Rho because a R135L-Rho-GFP fusion was previously shown to be mislocalized to the OPL when electroporated into WT rat rods (Hsu et al., 2015). While P23H-Rho is a class 2 autosomal dominant RP mutation that misfolds and causes ER retention, R135L-Rho is a class 3 RP mutation that causes mutant protein accumulation in the endocytic pathway (Aguilà et al., 2014; Athanasiou et al., 2018; Chuang et al., 2004). As controls, WT-hRho-RFP and WT-hRho-GFP AAVs were generated to express WT-Rho fusion proteins in WT rods. All AAV constructs include a C-terminal 1D4 tag fused in-frame after RFP or GFP.

3 weeks after subretinal AAV injection, retinas were screened with fluorescence microscopy to identify areas of high infection and no injection damage. Compared to WT-hRho-RFP, which predominantly localized correctly to the OS in transduced WT rods, P23H-hRho-RFP was mislocalized as bright puncta in the IS, encircling the nuclei throughout the ONL, and as distinct puncta in the OPL in transduced rods (Fig. 3A,B). This mislocalization pattern phenocopies the *P23H-RFP/+* mice and demonstrates that the P23H-hRho-RFP mislocalization is caused by the P23H-Rho mutation and not by non-autonomous effects in the disease-model transgenic mouse rods. WT-hRho-GFP also predominantly localized to the OS layer in transduced WT rods, while R135L-hRho-GFP accumulated as bright puncta at the IS/OS junction and mislocalized in a less bright but more consistent, diffuse pattern throughout the IS, ONL and OPL (Fig. 3C,D). Additionally, while WT Rho fusions were strictly localized in the OS in most WT transduced rods, there were some sporadic transduced rods with clear overexpression of WT-Rho fusion protein that mislocalized to the other photoreceptor cell layers (Fig. 3E,F).

**Figure 3.**
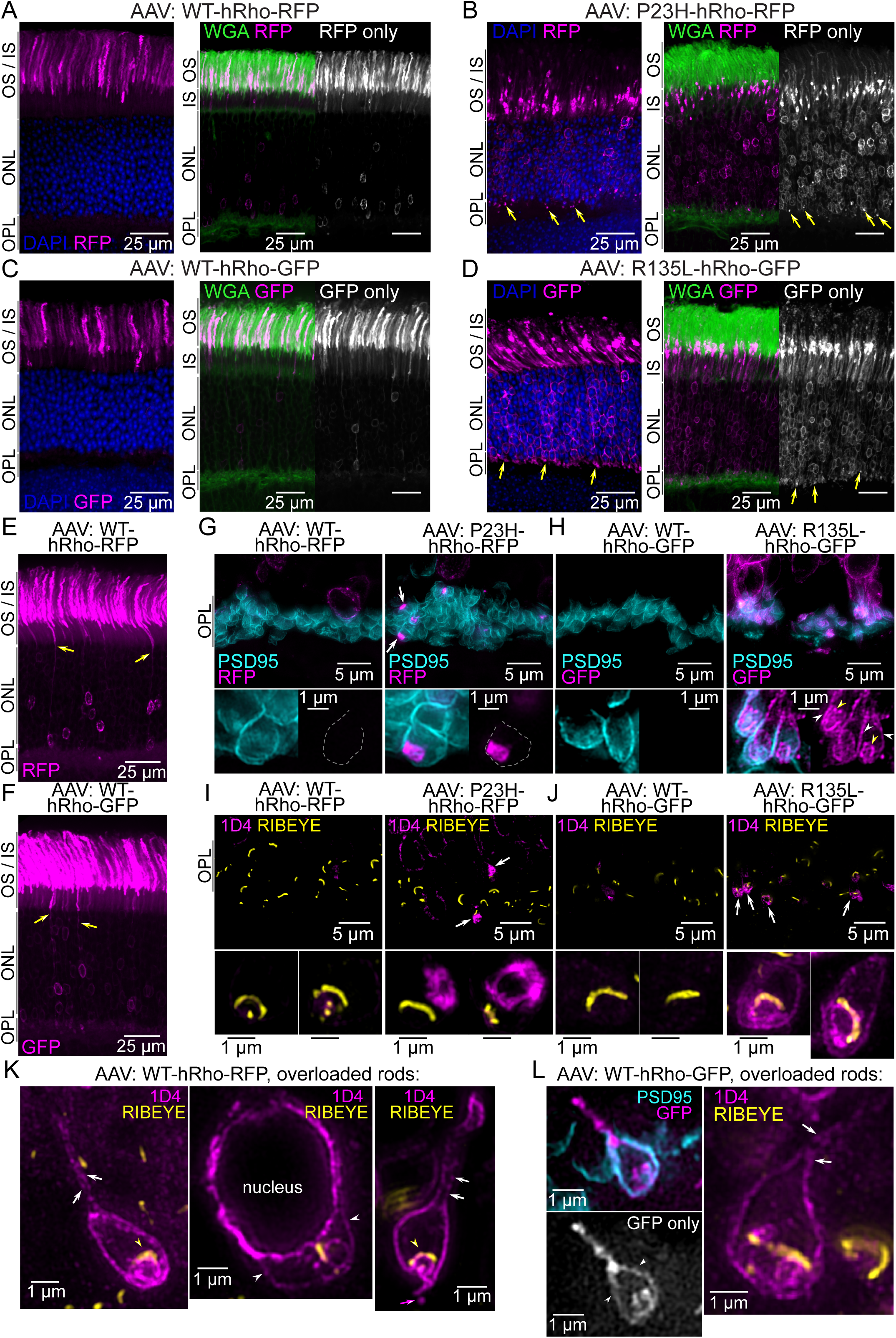
AAV overexpression of P23H-Rho and R135L-Rho in WT rods causes synaptic mislocalization. Confocal images of retinal sections from WT mice transduced with AAVs for rod-specific expression of the following Rho fusions: (A) WT-hRho-RFP, (B) P23H-hRho-RFP, (C) WT-hRho-GFP, and (D) R135L-hRho-GFP. Images on left side show Rho fusion localization (magenta) in transduced rods with DAPI (blue) labeling. Right images show co-labeling with WGA (green) as a marker for OS membranes. Grayscale images are the Rho-GFP/RFP fusion channels only. Yellow arrows = mutant Rho fusion OPL mislocalization. (E, F) AAV infected retinal sections with rods overexpressing (E) WT-hRho-RFP or (F) WT-hRho-EGFP (magenta, yellow arrows). (G, H) SIM z-projection images of the OPL from 3-micron retinal cryosections from the same AAV conditions as in (A-D). Rho fusion fluorescence is (magenta), and PSD95 immunolabeling (cyan) was used to identify rod spherules. Single spherule examples (bottom) of each AAV-driven Rho fusion are shown with PSD95 co-labeling (left) and the Rho-RFP/EGFP signal only (right). In (G), white arrows indicate P23H-hRho-RFP cytoplasmic aggregates, and dashed gray lines outline the PSD95+ plasma membrane of the magnified spherules of interest. In (H), R135L-hRho-GFP mislocalization at the spherule plasma membrane (white arrowheads) and internally (yellow arrowheads) are indicated. (I, J) SIM z-projection images from thin 1- micron sections of retinas from the same AAV conditions as in (A-D). These sections were co-immunolabeled for 1D4 (magenta) and RIBEYE (yellow). White arrows = mutant Rho fusion accumulates near the synaptic ribbon. Single spherule examples from each AAV condition are enlarged below. (K) SIM super-resolution images of WT-hRho-RFP overloaded rod spherules. 1D4+ WT-hRho-RFP (magenta) was localized along the plasma membrane of the axon and spherule (white arrows) and was colocalized with the RIBEYE synaptic ribbons (yellow arrowheads). In the far-right example, 1D4+ WT-hRho-RFP appears to bud off from the presynaptic spherule (magenta arrow). (L) SIM images of WT-hRho-GFP AAV overloaded rod spherules. WT-hRho-GFP colocalizes with the PSD95+ rod spherule plasma membrane (white arrowheads) and localizes at the rod axon plasma membrane (white arrows).

Next, AAV-transduced WT retinas were imaged with SIM to visualize the mislocalization of the mutant P23H-hRho and R135L-hRho fusion proteins in single rod spherules on a subcellular scale. First, cryosections of AAV-transduced retinas were used to preserve the Tag-RFP-T and EGFP fluorescence and were co-immunolabeled for PSD95 to label rod spherule plasma membranes for comparison to the RFP or GFP signal. In these samples, the mislocalized P23H-hRho-RFP in the OPL were bright aggregates, while no consistently strong signal above the background in the OPL was detected in control WT-hRho-RFP transduced retinas (Fig. 3G). In single spherules, the P23H-hRho-RFP aggregates were localized within the cytoplasm, which phenocopies the subcellular localization of mislocalized P23H-hRho-RFP aggregates in the OPL of *P23H-RFP/+* retinas. R135L-hRho-GFP mislocalization was also bright and apparent in the OPL of AAV-transduced WT retinas compared to WT-hRho-GFP, which was not detected in the OPL for most transduced areas (Fig. 3H). In single spherules, mislocalized R135L-hRho-GFP did not aggregate like P23H-Rho-RFP. R135L-hRho-GFP colocalized at the plasma membrane with PSD95 and internally in a pattern that suggests filling of the spherule plasma membrane surrounding the postsynaptic invaginations (Fig. 3H, yellow arrows).

For enhanced SIM resolution, we prepared thin plastic sections of AAV-transduced retinas. In this case, the Rho fusions were immunolabeled with the 1D4 antibody along with RIBEYE co-immunolabeling of the rod synaptic ribbons. In these sections, the mislocalized P23H-hRho-RFP aggregates were visualized as swirling patterns of aggregated mutant protein near the synaptic ribbon (Fig. 3I), again phenocopying the mislocalization pattern described in *P23H-RFP/+* retinas (Fig. 1F-G). R135L-hRho-GFP, on the other hand, was localized at the spherule plasma membrane and internally, partially colocalized with the synaptic ribbon (Fig. 3 J), again suggesting that R135L-hRho-GFP fills in the invaginating plasma membrane. Among all WT retinas infected with either WT-hRho fusion, we observed a few examples of 1D4 staining in the OPL in sporadic cells with dramatically overexpressed levels of WT-Rho. In these overloaded rod spherules, WT-hRho-RFP and WT-hRho-GFP were mislocalized along the axons and throughout the spherule plasma membrane, including within the plasma membrane surrounding the synaptic invaginations (Fig. 3K,L) in a pattern similar to R135L-hRho-GFP mislocalization.

### P23H-Rho-RFP mislocalization causes specific changes in the abundance of rod presynaptic proteins

Since the persistent accumulation of mutant P23H-Rho protein within large ER folds in the cytoplasm of *P23H-RFP/+* rod spherules caused no significant ultrastructural defects to the synaptic ribbons, we considered if the distension of the ER membranes in the spherules and cell bodies of *P23H-RFP/+* rods disrupted normal rod synaptic protein levels. In rod photoreceptor spherules, the ribbon is composed of structural proteins, including RIBEYE (Schmitz et al., 2000) and BASSOON (Bsn, (Dick et al., 2003)), and the ribbon is located just above the synaptic cleft where Cav1.4 L-type voltage-gated calcium channels (made of subunits Cav1.4α1-subunit and Cavβ2-/α2δ4 subunits) are localized and maintained by proteins like the active zone protein RIMS2 (Grabner et al., 2015; Lee et al., 2015). Rods also express cell-adhesion proteins that maintain their trans-synaptic connections with rod ON-type bipolar cells. These include ELFN1, which complexes with postsynaptic mGluR6 (Grm6) (Cao et al., 2015), and Dystrophin (Dmd), Dystroglycan (Dag1), and Pikachurin (Egflam1), which together complex with postsynaptic GPR179 (Furukawa et al., 2020; Omori et al., 2012; Orlandi et al., 2018; Sato et al., 2008) (Fig. 4A). These trans-synaptic proteins are aligned in the spherule plasma membrane surrounding invaginating ON-type bipolar cell dendrite tips, which was visualized with ELFN1 immunolabeling and SIM (Fig. S3A).

**Figure 4.**
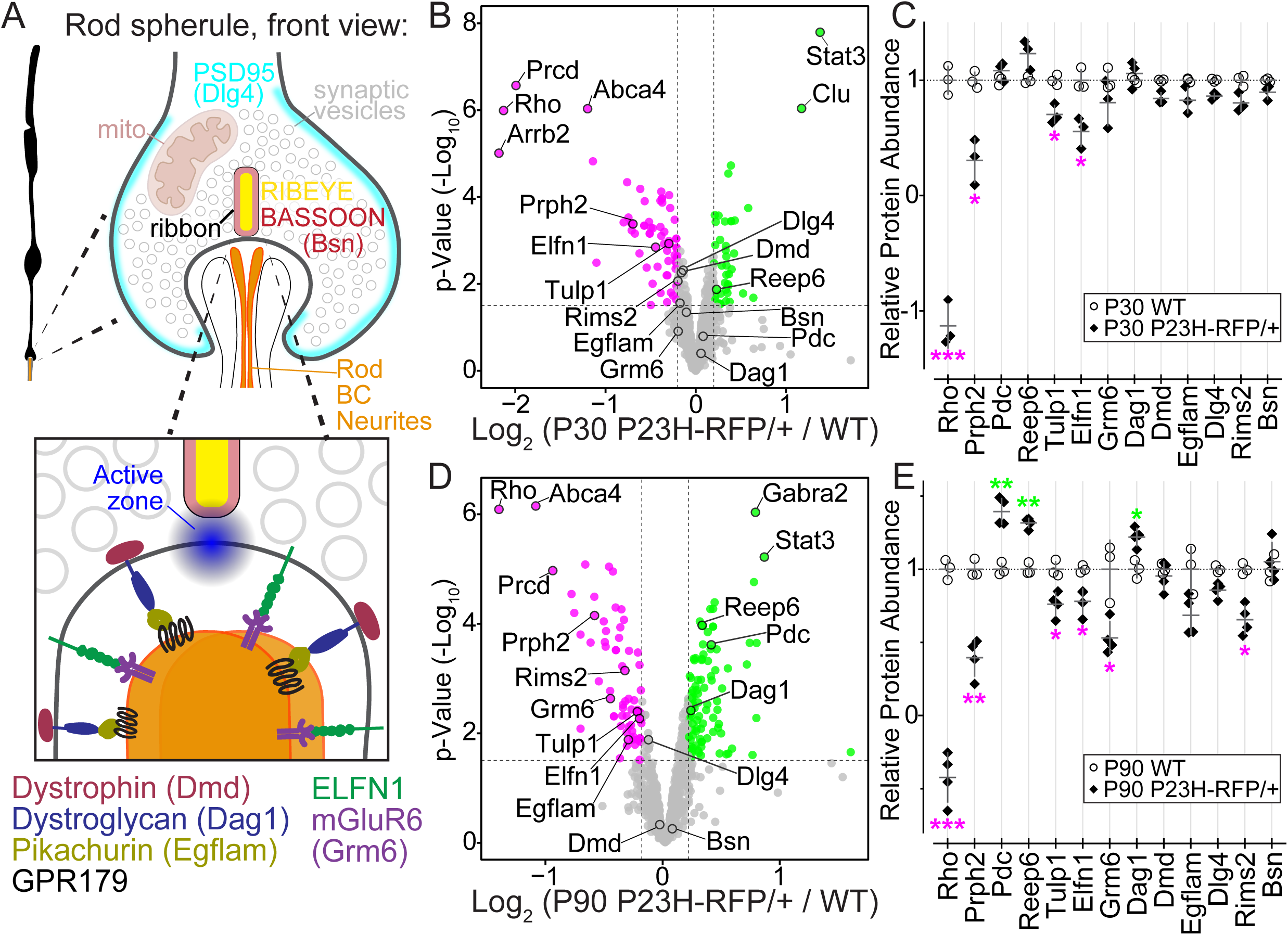
Photoreceptor and synaptic protein abundance changes are found in *P23H- RFP/+* retinas with TMT-MS proteomics. (A) Diagram of the front view of a rod spherule and a magnified view of the active zone and the synaptic cleft to highlight the approximate localizations of rod synaptic proteins including key trans-synaptic protein complexes. (B) Volcano plot of TMT-MS relative peptide abundances for select photoreceptor and synaptic proteins (based on Gene Ontology Cellular Component classification, see Methods) in age P30 *P23H-RFP/+* vs WT retinas. X axis = Log_2_ fold change values with a significance threshold of 0.2, Y-axis = p-values (-Log10) with a significance threshold of 1.5. Green points represent protein targets above the thresholds and magenta points are targets below the thresholds. Annotated protein names are based on FASTA gene names. (C) Linear scale graph of select relative protein abundances from the P30 TMT-MS data in Fig. S3B. WT (open circles) were normalized to 1 and superimposed with *P23H-RFP/+* values (black diamonds). Magenta asterisks = significant downregulation, and green asterisks = significant upregulation. (D) TMT-MS volcano plot comparing relative peptide abundances in age P90 *P23H-RFP/+* and WT retinas for the same protein list and plot parameters as in (B). (E) Linear scale graph of select relative protein abundances from the P90 TMT-MS data in Fig S3B with the same normalization and formatting as in (C).

To assess protein level differences between *P23H-RFP/+* and WT rods, tandem mass-tag mass spectrometry (TMT-MS) was performed on whole retina samples at ages P30 and P90. At both ages, there were significant peptide abundance changes in *P23H-RFP/+* retinas compared to age-matched WT controls (Fig. 4B,D, Fig. S3B). As expected, Rho peptides were significantly downregulated in *P23H-RFP/+* mice at both ages along with peptides for the OS-specific protein Prph2 and Tulp1, a trafficking regulator in rods localized in the IS, CC, and spherules (Hagstrom et al., 1999; Hagstrom et al., 2001) (Fig. 4B-E). Interestingly, peptide abundance for phosducin (Pdc), another OS protein, remain unchanged at P30 (Fig. 4B-C), but was significantly increased in P90 *P23H-RFP/+* retinas compared to age-matched WT controls (Fig. 4D,E). Additionally, in P90 *P23H-RFP/+* retinas, peptides for Reep6, an ER protein that was previously localized in the IS and OPL of mice (Agrawal et al., 2017), were significantly increased, while peptides for the rod synaptic proteins Elfn1, Rims2, as well as for Grm6 were significantly decreased (Fig.4E). Interestingly, peptide abundance for the rod trans-synaptic receptor Dag1 was significantly increased at P90 in *P23H-RFP/+* rods; however, there were no significant changes for its binding partner, Dystrophin (Dmd), at either age (Fig. 4D,E). TMT-MS peptide abundance differences for rod synaptic proteins such as Dystrophin may not have been detected due to their expression in other synapses of the inner retina which were present in our whole retina samples (Wersinger et al., 2011). Dystrophin protein isoforms (the Dp427, Dp260, and Dp140) and Dag1 protein levels were also not significantly changed in western blots from *P23H-RFP/+* and age-matched WT retinas (Fig. S3C).

Next, quantitative confocal microscopy was used to evaluate rod synaptic protein level changes specifically in the OPL of *P23H-RFP/+* and age-matched WT retinas to complement the whole retina TMT-MS data. In confocal fluorescent images, immunolabeled Dystrophin and ELFN1 localized as bright puncta in the OPL among the mislocalized P23H-Rho-RFP, while BASSOON (Bsn) immunolabeled the horseshoe-shaped synaptic ribbons (Fig. 5A-D). Dystrophin and Bsn were also localized in cone pedicle synapses (Fig. 5A, arrowheads), which are located proximally to rod spherules in the OPL (Moser et al., 2020). ELFN1 is specific to adult rod spherules (Cao et al., 2015; Cao et al., 2020), although we consistently observed an above-background ELFN1 immunofluorescence signal in the ONL of *P23H-RFP/+* retinas at P30 and P90 (Fig. 5C,D, Fig. S4A,B).

**Figure 5.**
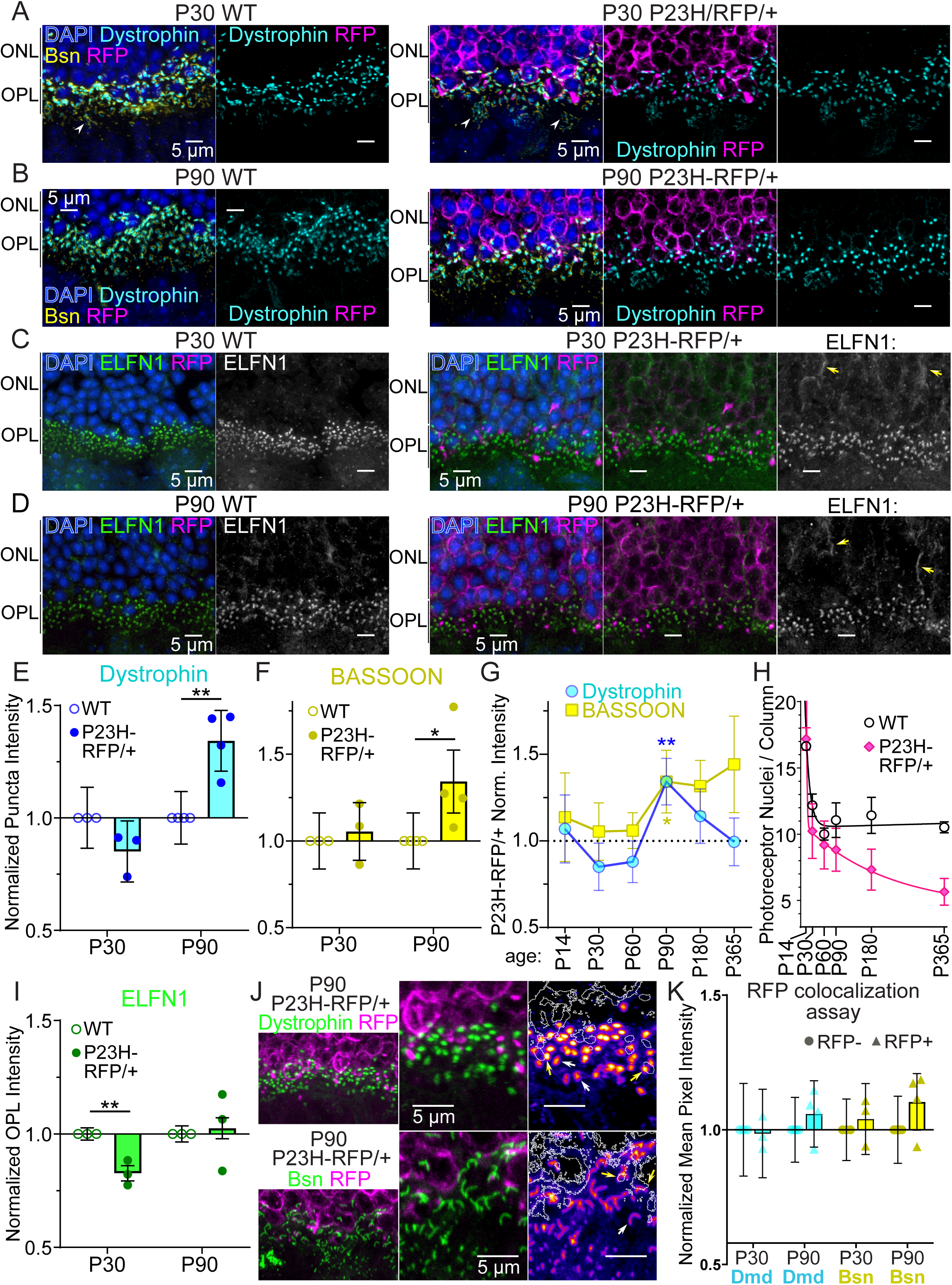
Quantitative confocal imaging analysis of synaptic protein abundance changes in *P23H-RFP/+* mice. (A, B) Example confocal z-projections from WT and *P23H-RFP/+* retinal cryosections at (A) P30 and (B) P90 focused on regions of the lower/proximal ONL and OPL with Dystrophin (cyan) and BASSOON (Bsn, yellow) immunolabeling and DAPI counterstaining (blue). RFP acquisition and intensity levels were matched between WT and *P23H-RFP/+* samples, and RFP fluorescence (magenta) was detectable only in the ONL and OPL of the *P23H-RFP/+* sections. (C, D) Confocal z-projections for WT and *P23H-RFP/+* retinal cryosections at (C) P30 and (D) P90 with ELFN1 immunolabeling (green) and DAPI nuclear staining (blue). RFP fluorescence (magenta) was again only detected in the *P23H-RFP/+* images. (E, F) Graphs of normalized puncta intensity values for (E) Dystrophin and (F) BASSOON from confocal images of WT and *P23H-RFP/+* retinas at P30 and P90. Graphs are aggregated data from replicate WT vs. *P23H-RFP/+* puncta intensity comparisons; for each comparison, and all replicates were aggregated. In the graphs, WT values are open circles and *P23H-RFP/+* are closed circles. (G) Time course plot of Dystrophin (blue) and Bassoon (yellow) normalized (Norm.) puncta intensity measurements from *P23H-RFP/+* retinas P30 and P90 data are the same aggregated replicates as in (E) and (F). (H) Time course plot of DAPI+ photoreceptor nuclei per column counted from both WT and *P23H-RFP/+* in confocal images analyzed throughout the puncta analyses in (E - G). Black circles = WT nuclei counts. Pink diamonds = *P23H-RFP/+* nuclei counts. Two phase decay curve fits were added in GraphPad Prism. (I) Graph of normalized ELFN1 OPL intensities between WT and *P23H-RFP/+* retinas at ages P30 and P90. The data were normalized and aggregated as in (E) and (F). (J) Example confocal images from *P23H-RFP/+* retinas depicting RFP fluorescence (magenta) colocalized with either Dystrophin (top, green) or BASSOON (Bsn, bottom, green) immunolabeling. In magnified views, on the right, Dystrophin and Bsn intensities are heat map pseudocolored and superimposed with the RFP signal as white outlines. White arrows indicate RFP+ Dystrophin/Bassoon colocalizations and yellow arrows indicate RFP- Dystrophin/Bassoon examples. (K) Graph of averaged puncta intensity values from the P90 *P23H-RFP/+* RFP colocalization assay for Dystrophin (cyan) BASSOON (yellow). The middle lines in each bar represent mean values.

For Dystrophin and Bsn, a single spherule puncta intensity analysis was performed using confocal imaging to evaluate protein level differences in *P23H-RFP/+* vs. WT rods at ages P30 and P90. At P30 there were no significant differences, but at P90, Dystrophin and Bsn levels were increased in *P23H-RFP/+* retinas (Fig 5E,F). Dystrophin and Bsn protein levels were also evaluated along a time course from ages P14 - P365, and P90 was the only time point in which there is a detectable difference among these proteins; however, there were consistently higher Bsn levels at older timepoints (Fig. 5G, Fig. S4C,D). DAPI+ nuclei were counted in all P14-P365 replicate mice in this analysis and plotted in Fig. 5H, and the rate of photoreceptor degeneration in *P23H-RFP/+* mice closely matches previous measurements (Robichaux et al., 2022). Interestingly, the P90 intensity increases for Dystrophin and Bsn correspond to the approximate timepoint where photoreceptor loss in *P23H-RFP/+* retinas plateaus. ELFN1 is rod-specific in the OPL; therefore, ELFN1 whole layer OPL immunofluorescence intensities were compared between *P23H-RFP/+* retinas and age-matched WT mice since there was no interfering cone signal. At P30, ELFN1 levels in the OPL were significantly reduced in *P23H-RFP/+* retinas but there was no difference at P90 (Fig. 5I).

Since the RFP+ signal in the *P23H-RFP/+* OPL confocal images were bright, easily detectable puncta, an additional analysis was performed comparing the intensities of Dystrophin and Bsn single spherule signals that were either colocalized with RFP puncta (RFP+) or not (RFP-) in the OPL of P30 and P90 of *P23H-RFP/+* retinas (Fig. 5J). There were no significant RFP+ vs. RFP- differences in the aggregated data (Fig S5E). This result indicates that synaptic proteins are disrupted in all *P23H-RFP/+* rods, possibly due to P23H-Rho mislocalization in the ER throughout all the inner rod compartments, not just due to ER aggregation in the spherules.

We next evaluated the above-background ELFN1 immunofluorescence in the ONL that was observed in *P23H-RFP/+* retinas (Fig. 5C,D, Fig. S4A,B) by quantifying ELFN1 and mGluR6 localization throughout the outer retina. In P30 *P23H-RFP/+* and age-matched WT retinas, ELFN1 and mGluR6 immunofluorescence were predominantly localized as overlapping puncta in the OPL, and ELFN1 signal was detected in the ONL of both genotypes (Fig. 6A,B and Fig. S5A,B). Upon closer analysis of the OPL in these co-labeled sections, there was a significant decrease in ELFN1 and mGluR6 spherule puncta labeling in *P23H-RFP/+* OPLs (Fig. 6C) but no evidence of mGluR6 mislocalization, which was previously shown in *Elfn1* knockout mice (Cao et al., 2015). Layer-specific pixel intensity measurements from these confocal images confirmed significant reductions in both ELFN1 and mGluR6 in the *P23H-RFP/+* OPL compared to WT controls (Fig. 6D, Fig. S5C). Interestingly, ELFN1 in ONL was not significantly different between *P23H-RFP/+* vs WT retinas; however, there was a significant increase in the ONL/OPL ratio of ELFN1 in *P23H-RFP/+* retinas (Fig. 6D), indicating an overall imbalance in the distribution of ELFN1 in *P23H-RFP/+* mutant rods.

**Figure 6.**
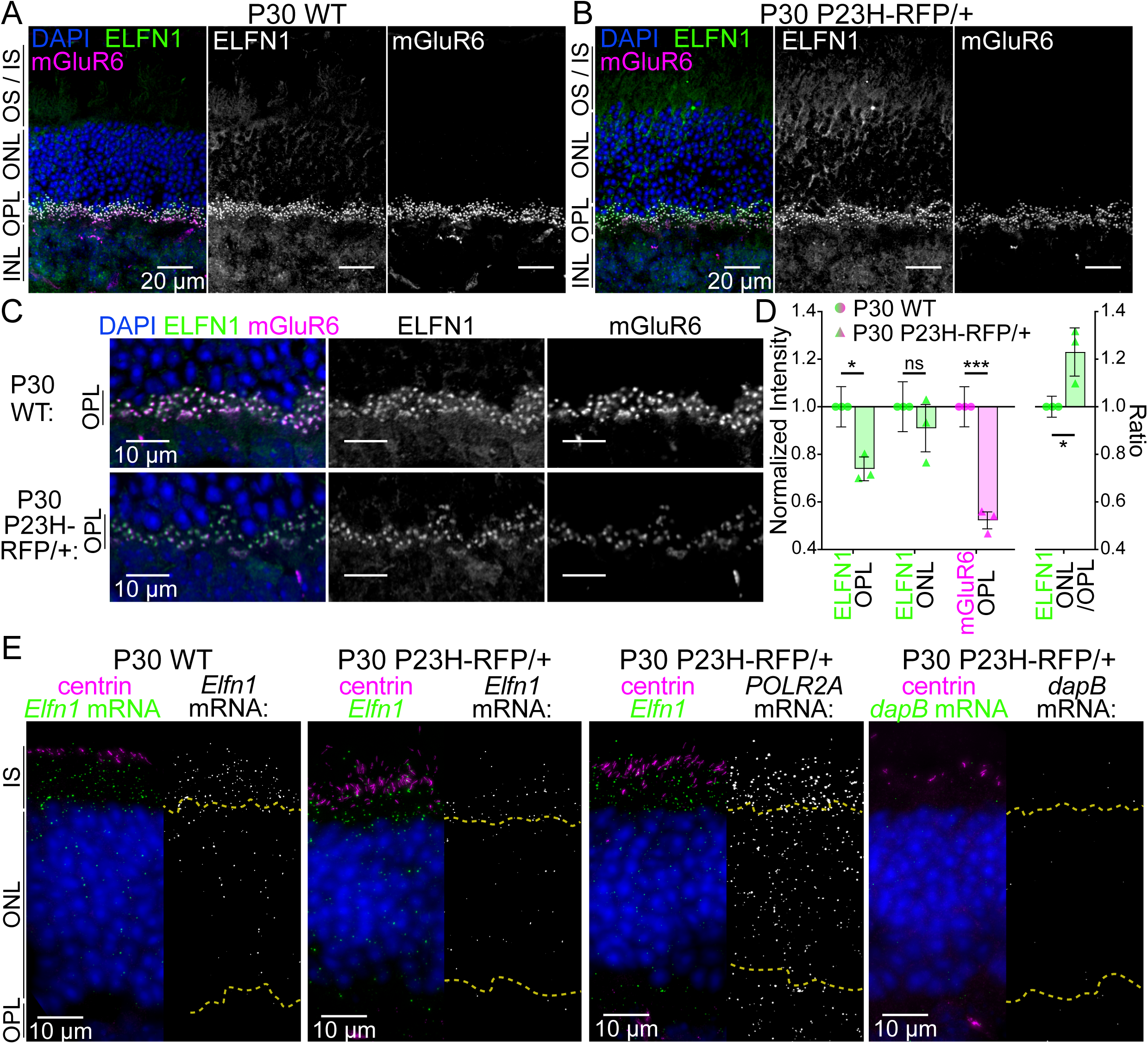
ELFN1 protein distribution is altered in *P23H-RFP/+* retinas. (A, B) Confocal z-projection images of (A) P30 WT and (B) P30 *P23H-RFP/+* retinal cryosections immunolabeled for ELFN1 (green) and mGluR6 (magenta) and counterstained for DAPI (blue). Grayscale images depict the ELFN1 and mGluR6 channels separately. (C) Magnified confocal images with the same labeling focused on smaller portions of the OPL. ELFN1 and mGluR6 staining, acquisition and intensity settings were matched throughout (A-C) between the P30 WT and P30 *P23H-RFP/+* sections. (D) Normalized intensity graph (left side) based on layer-specific intensity measurements for ELFN1 (green) and mGluR6 (magenta). Values were aggregated from replicate WT vs. P*23H-RFP/+* experiments; WT mean values are represented as circles and *P23H-RFP/+* values are triangles. On the right, ratios of ELFN1 intensity ONL/OPL intensities are plotted with the same parameters. (E) Example RNAscope SIM z-projections images for *Elfn1* mRNA (green) in P30 WT and P30 *P23H-RFP/+* retinas, alongside example SIM images probed for *POLR2A* (positive control) and *dapB* (negative control) mRNAs. Sections were co-immunolabeled for centrin (magenta) to label connecting cilia and counterstained with DAPI (blue). Grayscale images of mRNA puncta are shown to the right of the merged images. Yellow dotted lines = the IS:ONL and ONL:OPL boundaries based on the DAPI staining

Given this imbalance, *Elfn1* mRNA localization and abundance were evaluated using RNAScope fluorescence mRNA detection combined with SIM. With this method, *Elfn1* mRNAs were visualized as bright puncta localized throughout the IS and the ONL photoreceptor subcompartments in both P30 WT and P30 *P23H-RFP/+* retinas (Fig. 6E), demonstrating that *Elfn1* mRNA distribution was not grossly altered by the protein mislocalization and ER aggregation in *P23H-RFP/*+ rods at P30. Notably some *Elfn1* mRNA puncta were localized to the OPL, but these were sporadic and inconsistent. A positive control probe (*POLR2A*) targeting common housekeeping genes and a negative control probe (*dapB*) were analyzed for comparison. RNAScope puncta were counted to quantify any *Elfn1* mRNA abundance changes in the SIM data, and while *Elfn1* mRNA counts were significantly higher in the IS layer and significantly reduced in the OPL in *P23H-RFP/+* vs. WT P30 retinas in aggregated data (Fig. S5C), the counts were variable in the data among the *P23H-RFP/+* replicates (Fig. S5D). *POLR2A* positive control mRNA counts were significantly reduced in the distal ONL (dONL) and OPL in *P23H-RFP/+* retinas (Fig. S5E), indicating the possibility of a broader, cellular disruption to normal mRNA levels in *P23H-RFP/+* rods, which is further evidence that all *P23H-RFP/+* rods are impacted by protein aggregation throughout the ER.

### Rho mislocalization in rd10 RP mutant retinas does not induce rod presynaptic protein changes

For comparison with our *P23H-RFP/+* model, we next analyzed rd10 mice, an RP model known to exhibit synaptic mislocalization of Rho, in this case non-mutated WT Rho (Barhoum et al., 2008; Zhao et al., 2015). Rd10 mice contain a missense mutation in the beta subunit of phosphodiesterase-6 (PDE6β) that leads to rod degeneration beginning around P18 and peaking at ∼P20 (Gargini et al., 2007; Grossman et al., 2009; Jae et al., 2013; Wang et al., 2015; Zhao et al., 2015). Despite this early onset of degeneration, rd10 mutant rod ribbon synapses were shown to be morphologically normal at both P16 and P21 (Grossman et al., 2009). Here, confocal imaging of Rho immunofluorescence was used to examine rd10 retinas, which appeared relatively normal at P16, a pre-degeneration stage (Fig. 7A), but displayed degenerative morphology by P20 (Fig. 7B). Based on our imaging, rd10 retinas at P16 and P20 had Rho mislocalization to the IS, ONL, and OPL (Fig. 7A,B); however, the degree of Rho mislocalization in rd10 P16 retinas was not as robust as expected based on previous reports, nor was the mislocalization as striking as in the *P23H-RFP/+* retinas.

**Figure 7.**
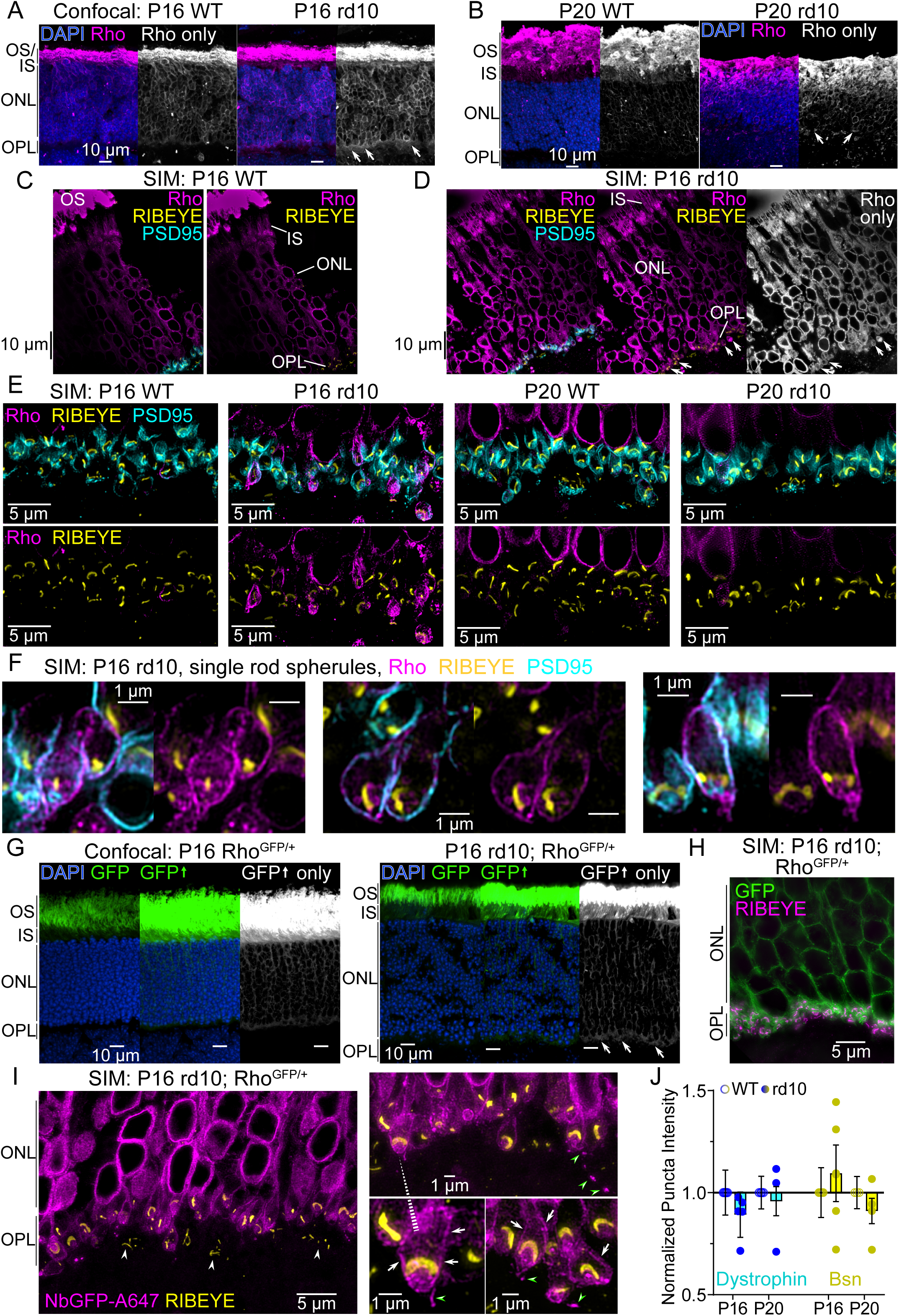
Rho mislocalization in rd10 RP mutant retinas does not alter rod synaptic protein levels. (A, B) Confocal z-projections of retinal cryosections from WT and rd10 mice at age P16 (A) or P20 (B). Sections were immunolabeled with the 4D2 Rho antibody (Rho, magenta) and counterstained with DAPI (blue). Rho only images are in grayscale. Mislocalized 4D2+ Rho signal is present in the rd10 OPLs (white arrows). (C) SIM images of a P16 WT retina immunolabeled for 4D2 (Rho, magenta), RIBEYE (yellow), and PSD95 (cyan). Rho fluorescence was detected throughout the OS, IS, and ONL but not in the OPL based on images with (left) and without (right) the PSD95 channel. (D) SIM image of a P16 rd10 retina labeled as in (C). Rho fluorescence was detected throughout the OS, IS, ONL, and also mislocalized in the OPL (white arrows) based on images with (left) and without (right) PSD95. The Rho channel only is in grayscale. 4D2 Rho immunolabeling, acquisition settings and intensity levels were matched between (C) and (D). (E) SIM images of the OPL in WT and rd10 mice at ages P16 and P20 with the same labeling and colors as in (C-D). 4D2+ Rho staining, acquisition and intensity settings were matched between all conditions, and in the WT OPL, the 4D2 signal is not consistently above background levels. In rd10 retinas at ages P16 and P20, mislocalized 4D2+ Rho labeling in the OPL is colocalized with PSD95 (top row) and surrounds the ribbons; Rho mislocalization in the P20 rd10 OPL is not as evident as at P16. (F) Magnified single rod spherule SIM images from P16 rd10 retinas with the same labeling as in (E). Mislocalized Rho colocalizes with PSD95 at the spherule plasma membrane (left images) and surrounds the ribbon (right images, no PSD95 channel). (G) Confocal z-projections of cryosections from Rho^GFP/+^ (left) and rd10; Rho^GFP/+^ (right) retinas depicting Rho-GFP (GFP, green) localization. Sections were counterstained with DAPI (blue). Grayscale images are GFP only with increased brightness to demonstrate Rho-GFP OPL mislocalization in rd10; Rho^GFP/+^ retinas (white arrows). (H) SIM image of a P16 rd10; Rho^GFP/+^ retinal cryosection immunolabeled for RIBEYE (magenta) to demonstrate that mislocalized Rho-GFP in the OPL overlaps with RIBEYE+ ribbons. (I) SIM images of thin resin sections of P16 rd10; Rho^GFP/+^ retinas that were co-immunolabeled with a GFP nanobody (NbGFP-A647, magenta) and for RIBEYE (yellow). NbGFP-A647 labeling was specific for Rho-GFP in these sections as cone ribbons are present (white arrowheads) with no surrounding NbGFP-A647 signal. In magnified views of rd10; Rho^GFP/+^ spherules, Rho-GFP is localized in the OPL along the spherule plasma membranes surrounding the ribbons (white arrows). Small Rho-GFP puncta were observed in the OPL extracellular space as if detached from the spherule membrane (green arrowheads). (J) Graph of aggregated normalized puncta intensities for Dystrophin (cyan) and BASSOON (Bsn, yellow) from age P16 and P20 WT (open circles) and rd10 (closed circles) retinas.

SIM imaging was used to examine the subcellular localization of Rho mislocalization in rd10 mice at P16 and P20. Retinas were immunolabeled with the 4D2 monoclonal Rho antibody (Hicks and Molday, 1986), a PSD95 fluorescent nanobody, and a RIBEYE antibody. While Rho immunofluorescence was detected in the IS and ONL of both P16 WT and rd10 retinas, the amount of Rho throughout the IS and ONL in P16 rd10 rods appeared higher. Rho was also clearly present in the rd10 OPL (Fig. 7C-E), which together indicates Rho mislocalization at this age. In the IS of P16 WT rods, we detected clear IS plasma membrane labeling (Fig. S6B), while in P16 rd10 rods, mislocalized Rho appeared to aggregate in the Golgi-rich IS myoid region and along the IS plasma membrane (Fig. S6C). In the ONL of both P16 WT and rd10 rods, Rho immunofluorescence surrounded the rod nuclei and also localized to the edges of ∼0.5-micron diameter projections corresponding to either the rod axons or the outer fibers that connect mouse rod ISs and cell bodies (Fig. S6D). In P20 rd10 retinas, Rho was also highly mislocalized in the IS compared to age matched WTs; however, OPL mislocalization was not evident at P20 (Fig. S6A) possibly due to advanced degeneration or labeling penetration issues. Notably in control P20 WT retinas, we were unable to detect Rho in the IS and most of the ONL, since P20 WT rods have more developed OSs that limit antibody penetration into the IS and ONL, as we previously described (Haggerty et al., 2024).

In SIM images focused on the OPL in age-matched WT and rd10 mice, Rho immunofluorescence signal was most abundant in P16 rd10 retinas, where mislocalized Rho labeled many teardrop-shaped spherules surrounding the RIBEYE+ rod synaptic ribbons, while in the P20 rd10 OPL, mislocalized Rho immunofluorescence was again less evident (Fig. 7E). In high magnification images of P16 rd10 rod spherules, mislocalized Rho was predominantly located along the spherule plasma membrane, colocalized with PSD95 (Fig. 7F, Fig. S6E). In some cases, the mislocalized Rho was also clearly localized along the axon that was continuous with the spherule plasma membrane, while in other cases, Rho appeared to be mislocalized internally, within the spherule cytoplasm and partially colocalized with the RIBEYE+ ribbon (Fig, S6E). However, it was not possible to distinguish if this signal represents Rho within the invaginating spherule plasma membrane, as seen for R135L-hRho-EGFP AAV transduced rods (Fig. 3H,J) and adult rods overexpressing WT hRho-RFP (Fig. 3K), or some other internal localization. Importantly, at this magnification, there was no detectable Rho immunofluorescence in P16 WT spherules (Fig. S6F).

To confirm this mislocalization pattern, rd10 mice were crossed with the *Rho-GFP-1D4* mice to generate a mouse homozygous for the rd10 mutation and heterozygous for Rho-GFP- 1D4 knockin fusion (rd10; Rho^GFP/+^). Using confocal imaging, Rho-GFP was mislocalized in the OPL in P16 rd10; Rho^GFP/+^ retinas compared to control Rho^GFP/+^ retinas (Fig 7G); however, as with Rho immunofluorescence staining in P16 rd10 retinas (Fig. 7A,B), the mislocalized Rho-GFP was clearly not accumulated or aggregated in the OPL. With SIM imaging of P16 rd10; Rho^GFP/+^ retinas, Rho-GFP fluorescence was first colocalized with the RIBEYE+ ribbons in the OPL (Fig. 7H), and then GFP nanobody staining was used to visualize Rho-GFP localization in single rod spherules. The Rho-GFP mislocalization pattern was again predominantly localized along the spherule plasma membrane, encasing the RIBEYE+ synaptic ribbons, with some potentially internal Rho-GFP that could be continuous with the Rho-GFP that was heavily localized in the ONL surrounding the rod nuclei (Fig. 7I). Additionally, we observed some Rho-GFP signal that branches away from the spherule into the extracellular matrix of the OPL (Fig. 7I, green arrowheads).

Finally, rd10 mice at ages P16 and P20 and age-matched WT controls were used to test for any Dystrophin and Bsn protein level changes in the OPL caused by Rho mislocalization. Upon quantitative confocal analysis, no significant differences were found for either protein between WT and rd10 mice at ages P16 and P20 (Fig. 7J), indicating that, unlike in *P23H- RFP/+* rods, these synaptic proteins are unaffected by Rho mislocalization in rd10 mice.

## Discussion

In this study, we found that P23H-Rho-RFP protein accumulated in ER membranes within presynaptic spherules of rod photoreceptor neurons from *P23H-RFP/+* RP mutant mice and that rod presynaptic protein levels were disrupted in these mutants. These results suggest the existence of a rod spherule ER-based secretory system that mediates proper synaptic protein trafficking and turnover and is sensitive to misfolded protein aggregation (Fig. 8). We also found three separate cases in which Rho mislocalized along the spherule plasma membrane: 1) in WT rods expressing non-aggregating but mislocalized RP mutant R135L- hRho-GFP (Fig. 3H,J), 2) in WT rods overloaded with WT-Rho-GFP/RFP fusion proteins (Fig. 3K,L), and 3) in rd10 RP mutant rods where a PDE6β mutation disrupts proper Rho trafficking which then overloads the rod plasma membrane (Fig. 7F,I). Thus, our findings provide new subcellular localization details of how different Rho mislocalization patterns differentially impact mouse rod presynaptic terminals.

**Figure 8.**
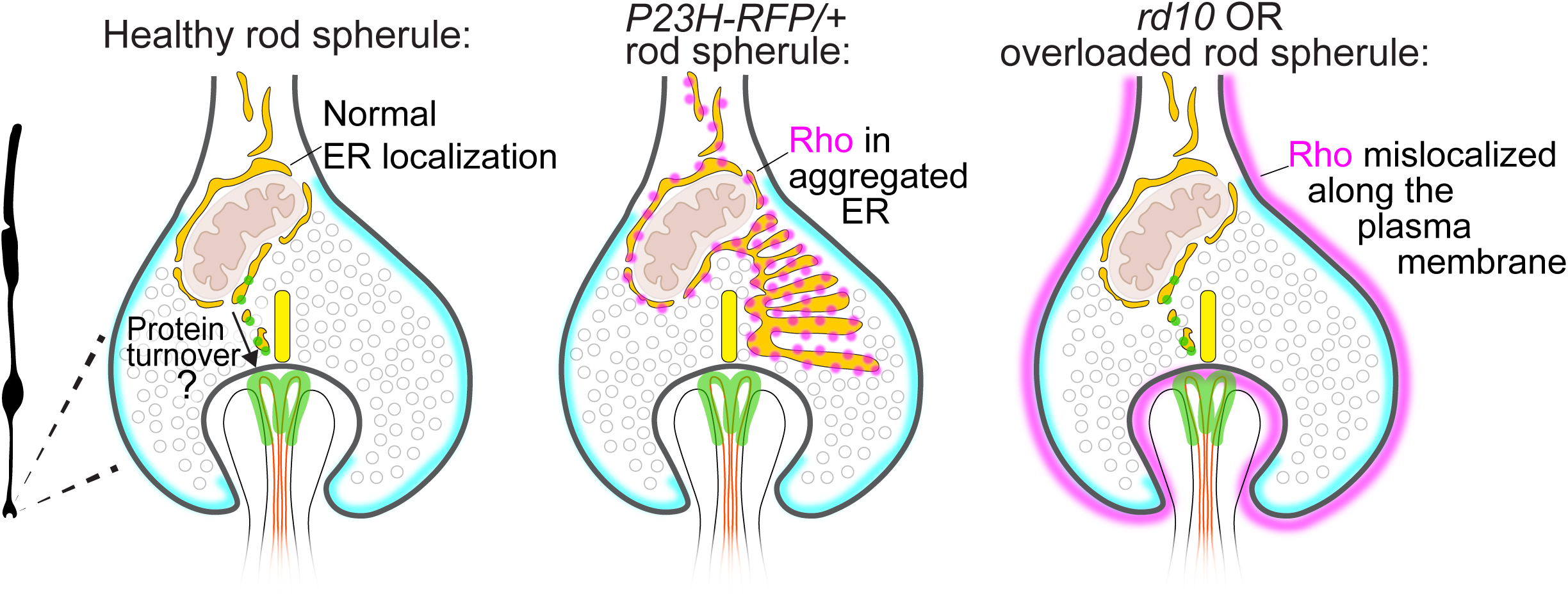
Diagram model of ER protein secretion in WT rod spherules and the impact of Rho mislocalization in mutant rod spherules. (A) Diagram of a WT rod spherule. The spherule plasma membrane is highlighted in cyan. ER (gold) wraps around the mitochondrion (tan) in the cytoplasmic space above the synaptic ribbon (yellow). Trans-synaptic cell adhesion proteins (green) align the synaptic cleft with the postsynaptic neurites (ON-type bipolar cells: orange; horizontal cells: black). These proteins (green) are potentially trafficked and turned over by an ER secretory pathway that extends to the rod spherule cytoplasm. (B) Diagram of a *P23H-RFP/+* rod spherule. Mutant Rho (magenta dots) aggregates in expanded ER (gold) in the cytoplasmic space of spherules, blocking the normal secretion of synaptic proteins. (C) Diagram of a rd10 rod spherule or a WT spherule with over-expressed WT-Rho fusion protein or mutant R135L-hRho-GFP protein. In these cases, Rho (magenta) mislocalizes along the spherule plasma membrane (cyan) such that the ER (gold) is putatively unaffected, and thus synaptic protein trafficking is normal.

While widespread P23H-Rho-RFP protein ER aggregation was clear throughout rods in mutant *P23H-RFP/+* retinas, ER aggregation was not originally apparent in the non-fusion *P23H-Rho/+* knockin mouse model (Sakami et al., 2011). However, a recent study using PROTEOSTAT, a dye that labels aggregated proteins, demonstrated mislocalized Rho aggregation surrounding photoreceptor cell bodies of the ONL in *P23H/+* mouse retinas peaking at age 3 weeks (Vasudevan et al., 2024). They also found that antigen retrieval was needed to detect aggregated Rho in *P23H-Rho/+* with the anti-1D4 Rho antibody, which may explain how P23H-Rho mislocalization was not previously detected with immunolabeling. Nevertheless, in the *P23H-RFP/+* mouse line used in this study, we confirm that ER aggregation of P23H-Rho-RFP extends throughout the rod inner compartments and into the OPL. Additionally, our newly generated *WT-RFP/+* mouse demonstrated that *P23H-RFP/+* mislocalization to synaptic aggregates are specifically triggered by the P23H-Rho mutation (Fig. 1B,C). Throughout this study, we examined the differences in Rho mislocalization between the *P23H-RFP/+* retinas and other models with SIM, which enabled us to clarify the extent of the ER network in mouse rods.

An inter-compartmental rod ER network that reaches the presynaptic spherules was previously described in amphibian rods (Mercurio and Holtzman, 1982). Furthermore, in salamander rods, ER-tracking dye and Ca^2+^ labeling demonstrated a soma-to-terminal ER-based intracellular Ca^2+^ network (Chen et al., 2014; Chen et al., 2015). Mouse rods are morphologically different: the spherules are significantly more segregated from the inner segment than in amphibian rods, such that mouse spherules have their own mitochondria. Additionally, most mouse rod spherules are R1-type, meaning that the spherules are also segregated from the cell body by a thin axon (Li et al., 2016). A study in mice used sarco/endoplasmic reticulum Ca^2+^-ATPase (SERCA) immunostaining established that presynaptic ER Ca^2+^ release helps sustain prolonged depolarizing conditions in rods (Babai et al., 2010). Our TEM imaging confirms these observations, as we visualized distinct ER-mitochondria associations between the ER and the mitochondrial outer membrane in *P23H- RFP/+* spherules (Fig. 2G,H). Other TEM studies noted ER membranes wrapped around the mitochondria in WT mouse rod spherules (Johnson et al., 2007; Ladman, 1958). Based on this association, the mouse spherule ER may form functional mitochondria-ER contacts that could participate in regulating proper glutamate release, lipid exchange, or Ca^2+^ signaling (Križaj, 2012; Tsuboi and Hirabayashi, 2021).

In other neurons, a peripheral ER network has been established as a long-range cellular network for protein trafficking and turnover (Tsuboi and Hirabayashi, 2021). In the axons of various neuronal cell types, a network of smooth ER was shown to form an “anastomosing” network of tubules (Broadwell and Cataldo, 1984; Konietzny et al., 2023; Lindsey and Ellisman, 1985) that wrapped around the mitochondria (Lee et al., 2018; Wu et al., 2017) and had distinct presynaptic ER cisternae branched off from the axonal ER network (Spacek and Harris, 1996; Wu et al., 2017). Based on our TEM imaging, we hypothesize that a similar ER network exists in mouse rod axons that wraps around the spherule mitochondrion and branches off smaller ER cisternae for protein exchange (Fig. 8). Since ER aggregation of P23H-hRho-RFP caused protein abundance changes, this presynaptic rod ER may be integrated with a cellular-wide protein synthesis and trafficking ER network that spans from the distal end of the IS to the spherules. Our finding of visible RFP+ aggregates in only ∼1 quarter of rod spherules from P30 and P90 *P23H-RFP/+* retinas suggests that the prolific Rho mislocalization throughout the IS of ONL in these mutant retinas (Fig. 1B,C) likely contributes to the widespread protein changes we measured. The distribution of smooth vs. rough ER in mouse rods remains an open question for future studies; the ER in our TEM data are densely stained and difficult to classify, but since we detected *Elfn1* and *POLR2A* mRNA throughout the IS, ONL and possibly the OPL (Fig. 6E), it is likely that rough ER is present in each of these compartments.

Among the synaptic protein abundances changes we discovered in *P23H-RFP/+* retinas, the depletion of ELFN1 in the OPL of *P23H-RFP/+* rods was the most significant change at age P30 (Fig. 5I, Fig. 6 A-D). In our TMT-MS data, *P23H-RFP/+* ELFN1 peptide abundances were also reduced at P30 and P90 (Fig. 4 C,E). Since ELFN1 localized as strings of fluorescence in the ONL from P30 *P23H-RFP/+* retinas throughout Fig. 5 and Fig. 6 resulting in a significantly imbalanced ELFN1 ONL/OPL distribution, we hypothesize that ELFN1 protein trafficking is specifically disrupted by P23H-hRho-RFP ER aggregation. Expectedly, this ELFN1 deficit coincided with significantly diminished postsynaptic mGluR6 puncta staining in the *P23H-RFP/+* OPL at P30 (Fig. 6 C-D). ELFN1 and mGluR6 interact to form a critical trans-synaptic complex between rod spherules and ON-type bipolar cell dendritic tips, as mGluR6 was also depleted in the OPL of *Elfn1* KO mice, which also had rod spherules with no invaginating ON-type bipolar cell dendrites (Cao et al., 2015).

Interestingly, at P90, *P23H-RFP/+* ELFN1 confocal immunofluorescence levels returned to WT levels (Fig. 5I), while Dystrophin and BASSOON levels were unchanged at P30 but then were significantly higher in P90 *P23H-RFP/+* retinas compared to WT controls (Fig. 5E-G, Fig. S4C,D). P90 *P23H-RFP/+* rods were described as having adaptive mechanisms that enabled some degree of normal Rho trafficking to the OS and cell survival (Robichaux et al., 2022). Upregulation of presynaptic proteins may be another such mechanism used by surviving P90 *P23H-RFP/+* rods to preserve essential rod synapses in response to long-term ER disruption. In support of this, a previous study using *P23H/+* heterozygous knockin mice demonstrated synaptic transcriptome increases at age 3 months (∼P90), which was attributed to homeostatic responses in rod bipolar cells to increase their sensitivity to rod input (Leinonen et al., 2020). Therefore, an adaptive strengthening of the rod-bipolar synapse may be a common phenotype in the middle stages of rod degeneration. Certain elements of the rod presynaptic machinery may be resilient to degenerating factors, since we found that the widespread ER aggregation in *P23H-RFP/+* rods caused no synaptic ribbon or vesicle docking ultrastructural defects despite the ER aggregates partially colocalizing ribbon in some SIM images (Fig. 1F,G).

In addition to ER aggregation in rods expressing P23H-hRho-RFP mutant protein, our visualization of mislocalized WT-Rho protein outlining the rod spherule plasma membrane either in WT rods overloaded with WT-Rho fusion protein (Fig. 3K,L) or in P16 rd10 rods (Fig. 7F,I), is informative about endogenous Rho trafficking dynamics. In our previous study, we used multiple labeling approaches with quantitative single molecule imaging to show that Rho is located throughout the IS plasma membrane in WT/non-diseased mouse rods (Haggerty et al., 2024). Thus, Rho that is overloaded in the IS may disperse throughout the entire rod plasma membrane and into the rod spherule plasma membrane including within the postsynaptic neurite invaginations. The mislocalized Rho in these overloaded regions may then be removed through endocytosis (Ropelewski and Imanishi, 2019) or exocytosis (Lewis et al., 2022), the latter of which we potentially observed here in the OPL in P16 rd10; RhoGFP/+ rods (Fig. 3K, Fig. 7I).

The differential impact of misfolded Rho protein ER aggregation vs. Rho overloading the rod plasma membrane is a crucial research question for future studies in order to better define the cellular status of rods in pre-degenerating disease states when they may be rescued by therapeutic intervention (Kunte et al., 2012; Leinonen et al., 2020). In this study we provide an in-depth examination of P23H-Rho-RFP ER aggregation in presynaptic rod spherules and evidence that some rod synaptic proteins are sensitive to these ER disruptions. An essential next step will be elucidating the trafficking mechanisms used for proteins like ELFN1 and Dystroglycan, which both require precise post-translational glycosylations to properly form trans-synaptic interactions (Park et al., 2020; Sato et al., 2008). Such future studies that elaborate the secretory system for rod synaptic proteins will provide much needed insight into subcellular disease mechanisms in rod neurons.

## Materials and Methods

### Animals

All WT mice (*Mus musculus*) were C57BL/6J. The *P23H-hRho-TagRFPt* (*P23H-RFP*) and the *Rho-GFP-1D4* mice were previously described in (Haggerty et al., 2024; Robichaux et al., 2022) and were also C57BL/6J. The rd10 mice (Jackson Laboratories) were also C57BL/6J and were crossed with *Rho-GFP-1D4* homozygotes to generate rd10;*Rho^GFP^* mice. *WT-hRho-TagRFP-T* (*WT-RFP*) mice were generated using CRISPR gene editing in the Genetically Engineered Murine Model Core Facility which is supported by the University of Virginia School of Medicine, Research Resource Identifiers (RRID):SCR_025473. The sgRNA (AGTACTGTGGGTACTCGAAGTGG) and HDR donor oligos (GCCACAGCCATGAATGGCACAGAAGGCCCTAACTTCTACGTGCCCTTCTCCAATGCGACG GGTGTGGTACGCAGT<u>CCC</u>TTCGAGTACCCACAGTACTACCTGGCTGAGCCATGGCAGTTC TCCATGCTGGCCGCCTACATGTTTCTGCTGAT) were developed for correcting the H23 mutant codon (CAC) to the WT P23 (CCC) codon in *P23H-RFP* mice. CRISPR reagents were injected into fertilized zygotes from *P23H-RFP* homozygous male and female mice for germline genome integration. Founder mice were then crossed to C57Bl/6J WT mice. F1 progeny were screened to confirm that the P23H mutation was corrected by sequencing genomic DNA. There was an unexpected silent change in the genome for residue S22 (AGC to TCA), but this did not change the coded amino acid sequence of hRho. We also validated the *WT-RFP* knockin with western blotting and fluorescence microscopy (Fig. 1 B-C, Fig. 1-1 A-B), and because validation was completed in the middle of this study, we continued using WT mice as controls for comparison to *P23H-RFP* mice throughout these experiments.

All mice were housed in 12 h light/dark conditions. Both sexes were used for experiments in the study unless otherwise noted. Mouse ages for imaging are denoted in the data. All experimental procedures using mice were approved by the Institutional Animal Care and Use Committee of West Virginia University (approval #2102040326).

### Antibodies and Labeling Reagents

The following primary antibodies were used in this study: anti-1D4 (Millipore Sigma, Cat# MAB5356, RRID:AB_2178961); anti-4D2 (Millipore Sigma, Cat# MABN15, RRID:AB_10807045); anti-Dystrophin (immunofluorescence) (Abcam, Cat# ab15277, RRID:AB_301813); anti-Dystrophin (western blotting) (Proteintech, Cat# 12715, RRID:AB_10640422); anti-Dystroglycan (Proteintech, Cat# 66735, RRID:AB_2934490); anti-RIBEYE (Synaptic Systems, Cat# 192103, RRID:AB_2086775); anti-BASSOON (Enzo, Cat# SAP7F407, RRID:AB_2313990); anti-TUBB5 (Millipore Sigma, Cat# MAB380, RRID:AB_2210541); anti-Sec61β (Cell Signaling Technology, Cat# D5Q1W); anti-Centrin, 20H5 (Millipore Sigma, Cat# 04-1624, RRID:AB_10563501); anti-ATPB2 (Proteintech, Cat# 22338, RRID:AB_2879077); FluoTag-X2 anti-PSD95-Alexa647 nanobody (NanoTag Biotechnologies, Cat# N3702-AF647, RRID:AB_2936216). The anti-ELFN1 polyclonal antibody was a gift provided by Dr. Kirill Martemyanov (University of Florida Scripps Institute). The anti-mGluR6 polyclonal antibody was a gift provided by Dr. Melina Agosto (Dalhousie University) (Agosto et al., 2021).

The following secondary antibodies were used in this study: F(ab’)2-goat anti-rabbit Alexa 488 IgG (Invitrogen, Cat# A11070, RRID:AB_2534114); F(ab’)2-goat anti-mouse Alexa 488 IgG (Invitrogen, Cat# A11017, RRID:AB_2534084); F(ab’)2-goat anti-rabbit Alexa 647 IgG (Invitrogen, Cat# A21246, RRID:AB_2535814); F(ab’)2-goat anti-mouse Alexa 647 IgG (Invitrogen, Cat# A21237, RRID:AB_2535806); F(ab’)2-goat anti-mouse Alexa 555 IgG (Invitrogen, Cat# A21425, RRID:AB_2535846); IRDye800CW goat anti-rabbit IgG (LI-COR, Cat# 925-32211, RRID:AB_2651127); IRDye800CW goat anti-mouse IgG (LI-COR, Cat# 925-32210, RRID:AB_2687825). Wheat germ agglutinin (WGA) staining was performed with WGA- CF640R (Biotium, Cat# 29206) and WGA-Alexa-CF488A (Biotium, Cat# 29022).

### Retinal Immunofluorescence

For immunofluorescence staining of mouse retinal cryosections for confocal microscopy, mouse eyes were enucleated, and the cornea, lens, and optic nerve were removed in ice cold 4% (wt/vol) paraformaldehyde (PFA) fixative. Eyecups were fixed for an additional 15 min at room temperature and transferred to a 30% sucrose in 1x phosphate-buffered saline (1xPBS) solution for 2 h on ice. Eyecups were further cryopreserved in a 1:1 mixture (OCT:30% sucrose) overnight at 4°C. Cryopreserved eyecups were frozen in optimal cutting temperature medium (OCT). 5 µm cryosections were cut on a Medical Equipment Source 1000+ cryostat, mounted onto Superfrost Plus slides (VWR Cat# 48311-701), and stored for less than 48 h at −80°C. For immunostaining, slides were warmed to room temperature prior to antigen retrieval in a 1X antigen retrieval solution (VWR, Cat# 103780-314) for 5 min at 80°C. Slides were equilibrated to room temperature, rinsed with 1xPBS, and incubated in an antibody blocking solution (10% Normal goat serum (NGS) + 0.1% Triton X-100 in 1xPBS) for 1 h at room temperature. Sections were incubated in 1–2 µg of primary antibodies diluted another blocking solution (2% NGS + 0.1% Triton X-100 in 1xPBS) at room temperature for 1 h. Sections were washed with PBS-T (0.1% Tween-20 in 1xPBS) 4 times for 5 min each prior to incubation with secondary antibodies diluted 1:500 in the same antibody blocking solution for 1 h at room temperature. Sections were washed in PBS-T and counterstained with 0.2 μg/ml 4′,6-diamidino-2-phenylindole (DAPI) (Thermo Fisher Cat# 62248) diluted in 1xPBS at room temperature for 20 min. Sections mounted with ProLong Glass Antifade Mountant (Thermo Fisher Scientific Cat# P36980).

For immunofluorescence labeling of whole retinas for structure illumination microscopy (SIM), mouse eyes were enucleated, and corneas were punctured and fixed in a 4% PFA + Ames’ media (Sigma Cat# A1420) fix solution for 15 min at room temperature. After the initial fixation, retinas were removed, bisected, and cut into trapezoid segments. Retinal segments were fixed in 4% PFA + Ames’ media for an additional 45 min at room temperature for a total 1 h fixation. Retinas used for anti-Sec61β ER immunolabeling were lightly fixed for only 5 min total and stained as whole retinas. Retinas were quenched in 100 mM glycine in 1X PBS for 30 min at 4°C and then incubated in SUPER block buffer (15% NGS, 5% bovine serum albumin (BSA) (Sigma, Cat# B6917) + 0.5% BSA-c (Aurion, VWR, Cat# 25557) + 2% fish skin gelatin (Sigma, Cat# G7041) + 0.05% saponin (Thermo Fisher, Cat# A1882022) + 1× protease inhibitor cocktail (GenDepot, Cat# P3100-005), in half dram vials (Electron Microscopy Sciences, Cat# 72630– 05) for 3 h at 4°C. Retinas were incubated with 5 µg of primary antibodies that were spiked into the block buffer for 3 full days, at 4°C with gentle agitation. A second dose of either Rho or Sec61β antibodies was added on the second day of primary antibody incubation to improve labeling. Retinas were washed 6 times with 2% NGS in Ames’ for 10 min each on ice prior to incubate with 4 µg of secondary antibodies in 2% NGS in Ames + 1X protease inhibitor cocktail for 12-16 h (overnight) at 4°C. Retinas were washed 6 times with 2% NGS in Ames’ 5 min each on ice and post-fixed in 2% PFA in 1xPBS for 30 min at room temperature with gentle agitation. Post-fixed retinas were then dehydrated with the following steps of pure ethanol diluted in water: 50%, 70%, 90%, 100%, 100%. Each ethanol step was performed at room temperature for 15 min with mild agitation. Following dehydration, retinas were embedded in Ultra Bed Low Viscosity Epoxy resin (Electron Microscopy Sciences, EMS Cat# 14310) using the following steps (all room temperature with gentle agitation): 1:3 resin to 100% ethanol for 2 h; 1:1 resin to 100% ethanol for 2 h; 3:1 resin to 100% for approximately 16 h (or overnight); 2 steps of full resin (no ethanol) 2 h each. Embedded retinas were then mounted in molds and cured for 24 h at 65°C. Resin blocks were trimmed and sectioned using glass knives on a Leica UCT Ultramicrotome to obtain 1 to 2 µm sections which were mounted on #1.5 glass coverslips with ProLong Glass mountant.

### RNAScope

Frozen 5 µm retinal sections were collected on Superfrost Plus slides as described above. Sections were dried for 1 h at −20°C and stored overnight at −80°C. The RNAScope Multiplex Fluorescent Reagent Kit v2 (Advanced Cell Diagnostics, ACD, Cat# 323110) was used for RNA detection, as follows. Sections were postfixed with 4% PFA in 1xPBS for 5 min at room temperature and then dehydrated in ethanol steps (50%, 70%, 100%, 100%) for 5 min each step. Sections were dried and treated with hydrogen peroxide for 10 min at room temperature. Sections were then incubated in boiling Co-Detection Target Retrieval (ACD, Ref#322000) for 2 min and then rinsed in water before incubation with 1 µg anti-centrin primary antibody (Millipore Sigma, Cat# 04-1624) overnight at 4°C. The following day, sections were fixed in a 10% neutral buffered formalin solution for 30 min at room temperature, washed, and incubated with Protease III (ACD, Cat# 322381) for 30 min in a 40°C hybridization oven. The RNA probes for *POLR2A*/positive control (ACD, Ref# 320881), *dapB*/negative control (ACD, Ref#320871), and *Elfn1*/Mm-Efln1 (ACD, Ref#449661) were added onto sections for a 2 h incubation at 40°C. Amplification steps were then subsequently performed per manufacturer instructions. Sections were then incubated with the HRP-C1 reagent (ACD, Cat# 323110) followed by the TSA vivid dye (ACD Cat# 323271, 1:25,000 diluted in TSA buffer (ACD, Ref# 322809)) for 30 min at 40°C for probe visualization. Finally, sections were incubated with secondary antibody (F(ab’)2-goat anti-mouse Alexa 647, diluted 1:500) for 1 h at room temperature. Sections were counterstained with DAPI for 30 seconds and mounted with ProLong Glass Antifade Mountant. Slides were imaged on a Nikon N-SIM E microscope system (see below), and z-projections (step size = 0.2 μm) were obtained for reconstruction. RNA probe SIM channels (488 nm) were processed for additional 3D deconvolution. Identical acquisition settings were used for each imaging field.

### Fluorescence Microscopy

Confocal microscopy was performed on either a Nikon C2 inverted confocal microscope, a Nikon AX inverted confocal microscope or a Nikon CrestV3 spinning disk microscope. Quantitative confocal imaging was performed either on the C2 and AX systems using Plan fluor 40x/1.30 NA (C2), Plan flour 40x/1.30 DIC H N2 (AX), or Plan Apo λ 100x/1.45 NA oil immersion objectives. 405 nm, 488 nm, 561 nm, and 640 nm laser lines were used on both systems. The CrestV3 spinning disk system was equipped with a Hamamatsu Fusion Gen III sCMOS camera, and a Plan Apo λD 60×/1.42 NA oil objective was used with Lumencor Celesta 405 nm, 477 nm, 546 nm, and 638 nm laser excitation. For all confocal imaging, identical acquisition settings were used between age-matched WT control and mutant sections, which were always mounted and immunolabeled on the same slide. Confocal images were acquired using Nikon NIS-Elements software and processed and analyzed using Fiji/ImageJ (Schindelin et al., 2009). SIM imaging was performed at room temperature as described in (Haggerty et al., 2024) using a Nikon N-SIM E microscope system equipped with a Hamamatsu Orca-Flash 4.0 camera and a SR HP Apochromat TIRF 100X, NA 1.49 oil immersion objective. Z-projections were obtained with 0.2 µm Z-steps (5-10 steps per image). SIM images were reconstructed using the NIS-Elements software and in some cases were additionally processed in NIS-Elements for 3D deconvolution using Automatic deconvolution mode. SIM images were processed and analyzed using Fiji/ImageJ.

### AAV Subretinal Injections

AAV constructs were designed and purchase from VectorBuilder. All AAV constructs contain a mouse rod specific MOPS500 promoter (Flannery et al., 1997). Human *RHO* coding sequences were tagged with an in-frame C-terminal TagRFP-T or EGFP. All AAVs contained an internal ribosome entry site (IRES) with a complimentary fluorescent tag (EGFP or mCherry) for visualization of AAV infection. AAVs were produced and packaged in the WVU Biochemistry and Molecular Medicine Virology Core. 3 adult WT mice were injected with each AAV. Subretinal injections were performed as in (Sechrest et al., 2024): prior to subretinal injection, mouse eyes were dilated with Tropi-Phen drops (Pine Pharmaceuticals). Mice were anesthetized using ketamine (80 mg/kg) and xylazine (10 mg/kg) in sterile 1xPBS via intramuscular injection. Fluorescein dye was added (0.1% final concentration) to AAVs for visualization. A 25-gauge needle was used to puncture the edge of the cornea. Transcorneal subretinal injections were performed by inserting a 33-gauge blunt end needle attached to a 5 μL Hamilton syringe containing 1 μL of AAV and injecting into the subretinal space. After injection, a Neomycin + Polymixin B Sulfates + Bacitracin Zinc ophthalmic ointment (Bausch & Lomb) was added to the eyes and antisedan (Orion Corporation) was intraperitoneally injected to reverse anesthesia.

21 days post-injection, mice were euthanized, and the injected eyes were enucleated. The corneas were punctured immersed in 4% PFA in Ames’ media. Eyes were fixed for 15 min before the cornea, lens, and optic nerve were removed. Eyecups were then embedded in 4% low melt agarose (Lonza, Cat# 50080). 150 µm vibratome sections were collected on a PELCO EasiSlicer vibratome and screened for AAV infection. Sections were stained for SIM by first quenching in a 100 mM glycine solution and blocking with 10% normal goat serum + 0.1% Triton X-100 in 1xPBS. 1-2 µg primary antibodies were spiked into the blocking solution and sections were probed for 12-16 h at 4°C with mild agitation. Sections were washed before incubation with secondary antibodies (diluted 1:500 in 1xPBS) for 2 h at room temperature. Sections were washed and post-fixed with 1% PFA prior to either sucrose cryopreservation and cryosectioning or ethanol dehydration and resin embedding as described above.

### Histology

For hematoxylin and eosin (H&E) staining and analysis, WT mice at age P30 (N=3) and *WT-RFP/+* mice at ages P30 and P180 (N=3, each age) were euthanized, and the eyes were enucleated and immediately incubated into Excalibur’s alcoholic Z-fix (Excalibur Pathology). Fixed eyes were sent to Excalibur Pathology, Inc. (Norman, OK) for paraffin sectioning and H&E staining. The H&E sections were imaged on a brightfield MIF Olympus Slide Scanner. 3 H&E retinal sections through the optic disk were used for each of the mice included in the analysis. Photoreceptor nuclei in the ONL were counted in a blinded analysis from an 80 µm wide central retina region located 500 µm for the optic disk for each section.

### TEM

Mouse eyes were enucleated, the anterior segments were removed, and the eyecups were immersion fixed in ice cold fixative (4% PFA + 2.5% glutaraldehyde diluted in 1xPBS, pH 7.4) for 12-16 h at 4°C with mild agitation. Retinas were dissected from the fixed eyecups and cut into four trapezoidal pieces. Retinas were rinsed with 100 mM cacodylate buffer (pH 7.4) 3 times and post fixed in 1% OsO4 + 1.5% K4[Fe(CN)6] × 3H2O in 100mM cacodylate buffer for 1 h at 4°C with mild agitation. Retinas were then washed three times in a wash solution (100 mM cacodylate buffer + 50 mM Na-maleate, pH 5.0, in water) on ice, 5 min each step. Then, retinas were incubated in 2% uranyl acetate (UA) in 50 mM Na-maleate for 3 h at 4°C with mild agitation. Retinas were washed with 50 mM Na-maleate 3 times and water 3 times on ice for 5 min each step before dehydration with increasing concentrations of ethanol (50%, 70%, 90%, 100%, 100%) and 2 100% acetone steps at room temperature for 15 min each step. Dehydrated retinas were resin-embedded in increasing concentrations of Eponate 12 resin (Ted Pella Inc., Cat#18010) in room temperature/mild agitation stages, as follows: 1:1 resin to acetone overnight; 3:1 resin to acetone 2 h; 2 full resin steps (no acetone), 2 h each. Embedded retinas were transferred to molds and cured at 65°C for 48 hours. Ultrathin resin sections (70 nm) were cute on a Leica UCT ultramicrotome using a Diatome Ultra 45° diamond knife and collected on copper grids (Electron Microscopy Sciences, Cat# G100-Cu). Copper grids were post stained with a 1.2% UA solution and a 3% lead citrate solution (Electron Microscopy Sciences Cat#22410) for 4 min each. Grids were imaged on either a Joel JEM 1010 transmission electron microscope or a Joel 1400 transmission electron microscope.

### TMT-MS

Whole mouse retinas were dissected for tandem mass tag mass spectrometry (TMT-MS) in sterile 1xPBS, and the ciliary bodies were removed. Retinas from both eyes of each mouse were combined and flash-frozen on dry ice. For P30 analysis, 3 WT and 3 *P23H-RFP/+* males were used. For P90 analysis, 3 WT female and 4 *P23H-RFP/+* female mice were used. Frozen retinal samples were sent to IDEA National Resource for Quantitative Proteomics (Little Rock, AR) for lysis, trypsin digestion, TMT labeling and Orbitrap Eclipse MS acquisition. Database analysis, quality control, normalization, fold change calculations and statistical testing (described below) were performed by IDEA. All acquired TMT-MS data is available in Table S1.

### Western Blotting

3 WT and 3 *P23H-RFP/+* mice were used for age P30 and P90 western blot analyses. Dissected retinas were flash frozen on dry ice for at least 10 min, resuspended in 100 µl of 1% Triton X-100 + 1X protease inhibitor (Thermo Fisher, Cat#A32955) in 1xPBS and lysed by sonication. Lysed samples were cleared with centrifugation, and 11 µl of each sample was combined with 11 µl of urea sample buffer (6 M urea + 140 mM SDS + 0.03% bromophenol blue + 360 mM BME in 0.125 M Tris (pH 6.8)) and heated for 5 min at 95°C. Samples were loaded into a Novex Tris-glycine mini 4-12% gel (Thermo Fisher Cat# XP04120) for SDS-PAGE. Gels were transferred onto Immobilon-FL Transfer Membrane polyvinylidene difluoride (PVDF) (pore size: 0.45 μm) (LI-COR Cat# 92760001) in Tris-Glycine Transfer Buffer (Bio-Rad Cat# 1610771). Blots were blocked with Intercept Blocking Buffer (LI-COR, Cat# 927–6000) for 1 h and then washed 3 times with PBS-T 5 min each. Primary antibodies at 1:500 to 1:20,000 dilutions in PBS-T were added to the blots for 2 h probing at room temperature. Blots were washed and secondary antibodies (diluted 1:50,000 in PBS-T) were added for 1 h probing at room temperature. Blots were imaged on an Amersham Typhoon scanner (GE).

### Deglycosylation Assay

Deglycosylation assays were performed as in Haggerty et al. (2024): dissected retinas were flash frozen at −80°C for 10 minutes and then lysed in 100 µL of RIPA buffer (Alfa Aesar, Cat# J63306) + protease inhibitor cocktail (GenDepot Cat# P3100-001). Samples were cleared with centrifugation and the cleared supernatant was used for the assay. Lysate was mixed with deglycosylation buffer containing PNGase F (New England Biolabs, Cat# P6044) and incubated for 10 min at 37°C. Protein deglycosylation mix II (New England Biolabs, Cat# P6044S) was added to the treated tubes and buffer only was added to the control tubes. All samples were then incubated for 1 h at 37°C. Samples were cooled on ice for up to 10 mins and mixed with urea sample buffer for a 1:1 mixture and loaded onto IVGN Novex WW 10-20% Tris-Glycine gels (Fisher Scientific, Cat# 89238-778) for western blotting, as described above.

### Image Processing and Analysis

For confocal image puncta analyses (Fig. 5E-F, Fig. S4C-D; Fig. 7J), channels of interest were cropped a width of 80 µm and coded for a blinded analysis. For Dystrophin and BASSOON, integrated densities of 10 individual puncta were measured for each of 4 images per mouse included in the analysis (40 puncta per mouse total). An equal number of mean background measurements were taken for each image. The average mean background was multiplied by the puncta areas and these values was used for background subtraction. For the ELFN1 OPL particle analysis, the OPL was selected based on DAPI staining and thresholding was used to collect the foreground ELFN1 puncta signal and background for each image. Thresholding values were identical for all images and conditions. For ELFN1 and mGluR6 ONL vs OPL intensity measurements (Fig. 6 A-D, Fig. S5C), mean intensity values were measured from ONL and OPL regions, which were selected based on the DAPI channel. For the RFP puncta analysis, channels were selected as before, and puncta were preselected and coded based on their association (RFP+) or lack of association (RFP-) with RFP signal for blinded integrated density measurements. RNAscope puncta were counted from SIM z-projections by selecting IS, ONL and OPL regions based on the centrin immunostaining and DAPI channels. In SIM images, the distal ONL (dONL) was distinguished in images focused on the IS region from the proximal ONL (pONL) in images focused on the OPL. Analyze Particles in FIJI/ImageJ was used for puncta counting in each region, and the same thresholding values were used for all images. For TEM, images that contained a clear ribbon in the front-view, rod-like orientation were selected for a blinded analysis based on the parameters described in (Kesharwani et al., 2021). The ribbon height was manually measured from the anchoring site to the ribbon tip, and the synaptic vesicles were manually counted and considered ribbon-associated if they overlapped or had clear tethers to the ribbon. For western blotting, the intensities of the bands from the blot scan images were measured using Fiji/ImageJ. Background measurements were also taken from the same area as the measured bands for background correction. Labeling on images is as follows: OS = outer segment, IS = inner segment, ONL = outer nuclear layer, OPL = outer plexiform layer. Scalebars match adjacent images when not labeled.

### Experimental Design and Statistical Analysis

Specific experimental design details, such as number of mice and cells examined, are included in the Results text or the figure legends. All confocal analyses were performed with matching retinal sections between experimental and WT control conditions. Experimental datasets were directly compared to the matching WT data, and thus all data were normalized for aggregation so that WT mean values = 1. To statistically compare the aggregated data, standard deviations were propagated to determine the relative error, and the propagated standard deviations are represented as error bars in each of the aggregated data graphs. For TMT-MS data, volcano plots were generated using VolcaNoseR (Goedhart and Luijsterburg, 2020), and gProfiler was used to classify proteins based on the Gene Ontology Cell Compartment/GO-CC terms: “photoreceptor outer segment,” “synapse,” “photoreceptor inner segment,” and “photoreceptor connecting cilium.” TMT-MS data were statistically compared with a differential abundance analysis and moderated t-tests to account for protein variability, distribution, and abundance. For each comparison, P-values and false discovery adjusted P-values were calculated. All graphs were generated using GraphPad Prism and all statistical testing was performed in either GraphPad Prism or GraphPad Quickcalcs. WT data were normalized to a value of 1, and mutant data was then normalized to this value. In all graphs: bars = aggregate mean aggregate values, error bars = standard deviations after error propagation unless stated otherwise. Unless specified otherwise, significance was determined using unpaired T-tests with error bars representing standard error. Asterisks indicate the following adjusted p-values: * P <0.055, ** P<0.01, *** P<0.001. All original data corresponding to the graphs are openly available via Mendeley Data at DOI: 10.17632/w4d49nkm5n.1.

## Acknowledgements

The authors thank Dr. Abigail Moye and Dr. Melina Agosto for feedback during manuscript preparation.

## Author contributions

Conceptualization: S.L.T., M.A.R.; Methodology: S.L.T., M.A.R.; Validation: S.L.T.; Formal analysis: S.L.T., M.A.R.; Investigation: S.L.T., S.M.C., M.H., E.R.S., W.D., M.A.R.; Resources: W.D., M.A.R.; Writing – original draft: S.L.T.; Writing – reviewing & editing: S.L.T., M.H., W.D., M.A.R.; Supervision: W.D., M.A.R.; Project administration: M.A.R.; Funding acquisition: W.D., M.A.R.

## Data and resource availability

All original data corresponding to the figure graphs are openly available via Mendeley Data at DOI: 10.17632/w4d49nkm5n.1. TMT-MS data is available in Table S1. All other relevant data and resources can be found within the article and the supplemental figures.

## DEI

Proper diversity, equity, and inclusion practices were used throughout the study to ensure equitable opportunities for those involved.

---

**Supplemental Figure 1.** (A) Western blots of WT (age P77), *Rho-GFP-1D4/+* (abb: *WT-GFP/+*, age P61), and *WT-RFP/+* (age P58) retinal lysates; 2% of the total volume from 1 mouse retina was used for each lane. In *WT-GFP/+* and *WT-RFP/+* lanes, 60-65 kDa Rho-C-1D4-positive bands corresponding to the Rho-GFP/RFP fusions are larger than the ∼35 kDa endogenous monomer Rho protein bands that is present in all lanes. Magenta arrow = the band corresponding to WT-hRho-RFP. (B) Western blots of WT (age P70), *WT-GFP/+* (age P67) and *WT-RFP/+* (age P58) retinal lysates after treatment with PNGase F or buffer only. Rho protein deglycosylation shifts to lower molecular weights were evident with 1D4 (Rho-C-1D4) immunolabeling, including shifts in the WT-hRho-RFP band (magenta arrow). Na, K ATPase beta 2 (ATP1B2) deglycosylation serves as the control (“Deglycos. control”). (C) H&E stained central retina example sections from P30 WT and P30 *WT-RFP/+* retinas. In the adjacent graph, ONL density values (# of photoreceptor nuclei in an 80 µm central retina region) are plotted for P30 WT, P30 *WT-RFP/+* and P180 *WT-RFP/+*. Points = measurements from replicate mice (N=3 each condition). Bars = mean values. (D) DAPI+ nuclei per ONL column values from confocal images like in Fig. 1 A-B were plotted for *WT-RFP/+* and *P23H-RFP/+* mice at ages P30 and P90. Points = mean values. (E) Original P30 WT SIM images without 3D deconvolution corresponding to Figure 1D. (F) Original P30 P23H-RFP/+ SIM images without 3D deconvolution corresponding to Figure 1E. (G) Original single spherule SIM images without 3D deconvolution corresponding to Figure 1F. (H) Original SIM images without 3D deconvolution corresponding to the images in Figure 1G. (I) Additional P90 P23H-RFP/+ single spherule examples (SIM and SIM + 3D Decon). White arrows = 1D4 (magenta) aggregates localized in the *P23H-RFP/+* OPL. White asterisks = gaps in the aggregated 1D4 fluorescence in P23H- RFP/+ spherules. (J) SIM and SIM + 3D Decon images of WT OPL at age P90. Some 1D4 signal can be observed in the ONL(white arrow) but not in the OPL. Throughout the figure scale bars match adjacent images when not labeled.

**Supplemental Figure 2.** (A) TEM images of P30 WT spherules (magenta asterisks = mitochondria, green arrowheads = endocytosed vesicles). ER-like membranes (orange arrowhead) are located near the mitochondria. (B) TEM images of P30 *P23H-RFP/+* rod spherules annotated as in (A). Denser ER membranes are observed in these mutant spherules (orange arrowheads) (C) A P30 WT R1-type rod spherule, in which ER (orange arrowheads) is observed in the axon and surrounding the spherule mitochondrion (magenta asterisk). (D) TEM image of an R2 spherule in a P90 WT retina depicting ER-like membranes (orange arrowhead) extending from around the nucleus and into the spherule of the cytoplasm.

**Supplemental Figure 3.** (A) SIM super-resolution image of the OPL of a P30 WT retina. The rod synaptic protein ELFN1 (cyan) is typically located beneath the synaptic ribbon (yellow). Gaps in a single ELFN1 fluorescent punctum suggest the shape of invaginating post-synaptic neurites (dotted white lines). (B) Table of normalized TMT-MS values for select proteins, including the proteins in Fig. 4 B and D. Dag1 = Dystroglycan. (C) Western blot analysis for Dystrophin isoforms in WT and *P23H-RFP/+* whole retinas at ages P30 and P90. Each lane represents a retinal lysate sample from a separate mouse. Colored brackets on the blots correspond to the colored bars in the graphs for densitometry intensity quantification. All intensities are plotted as normalized to corresponding tubulin band intensities (Tubb5, bottom blots).

**Supplemental Figure 4.** (A, B) Confocal z-projections of replicate WT (left) and *P23H-RFP/+* (right) retinas immunolabeled for ELFN1 (green) at ages (A) P30 or (B) P90. ELFN1 and RFP (magenta) levels were matched between WT and *P23H-RFP/+* images. Yellow arrows = strings of ELFN1 in the *P23H-RFP/+* ONL. The ELFN1 only channel is shown in grayscale. (C, D) Graphs corresponding to the data in Fig. 5 G. (C) Time course analysis of Dystrophin (cyan) normalized intensities ranging from age P14 to P365 in WT (open circle) and *P23H-RFP/+* (closed circle) retinas. (D) Time course analysis of BASSOON (yellow) normalized intensities ranging from age P14 to P365 in WT (open circle) and *P23H-RFP/+* (closed circle) retinas.

**Supplemental Figure 5.** (A, B) Confocal z-projection images of replicates for (A) P30 WT and (B) P30 P23H-RFP/+ retinal cryosections immunolabeled for ELFN1 (green), mGluR6 (magenta), and DAPI (blue). Merged images are adjacent to grayscale images of the separated ELFN1 and mGluR6 channels. (C) Graph depicting aggregated and normalized *Elfn1* mRNA particles per area (μm^2^) from the RNAScope examples data in Fig. 6 E. *Elfn1* counts are graphed for the IS, dONL, pONL, and OPL layers from P30 WT (circles) and P30 P23H-RFP/+ (triangles) retinas. Values are from replicate WT vs *P23H-RFP/+* comparisons and all data were normalized to WT mean = 1. (D) Graph of the de-aggregated, normalized Elfn1 mRNA particles per area data corresponding to Fig. S5C. (E) Graph of replicate, normalized mRNA particles per area values for the control RNAscope probes in Fig. 6E.

**Supplemental Figure 6.** (A) SIM super-resolution example images of P20 WT (left) and rd10 (right) retinas. 4D2+ Rho signal (magenta) is localized in WT rods at the bottom of the ONL near the spherules (cyan) and ribbons (yellow), but not in the OPL. Faint 4D2 staining in rd10 rods is detectable in the OPL. (B) 4D2+ Rho fluorescence in a SIM super-resolution image of a P16 WT retina. Rho is located along the IS plasma membrane (yellow arrows). (C) 4D2+ Rho fluorescence in SIM super-resolution example images of P16 rd10 retinas demonstrates Rho localization at the IS plasma membrane and myoid region. (D) More 4D2+ Rho fluorescence in SIM super-resolution images from P16 WT and rd10 retinas focused on the ONL. Here, Rho is localized along the membranes of rod outer fibers/axons. (E) SIM super-resolution single spherule example images from P16 rd10 retinas. 4D2+ Rho (magenta) is colocalized with PSD95 (cyan) at the spherule plasma membrane and along rod axons (white arrows). (F) SIM super-resolution single spherule images from P16 WT retinas. 4D2+ Rho staining, acquisition and intensities were matched to all the examples in (E), (F), and Fig. 7 F. There is only sporadic, faint Rho (magenta) signal in some WT examples.

