## Supplementary material for "P23H rhodopsin aggregation in the ER causes synaptic protein imbalance in rod photoreceptors": Figure S

Supplemental Figure 1

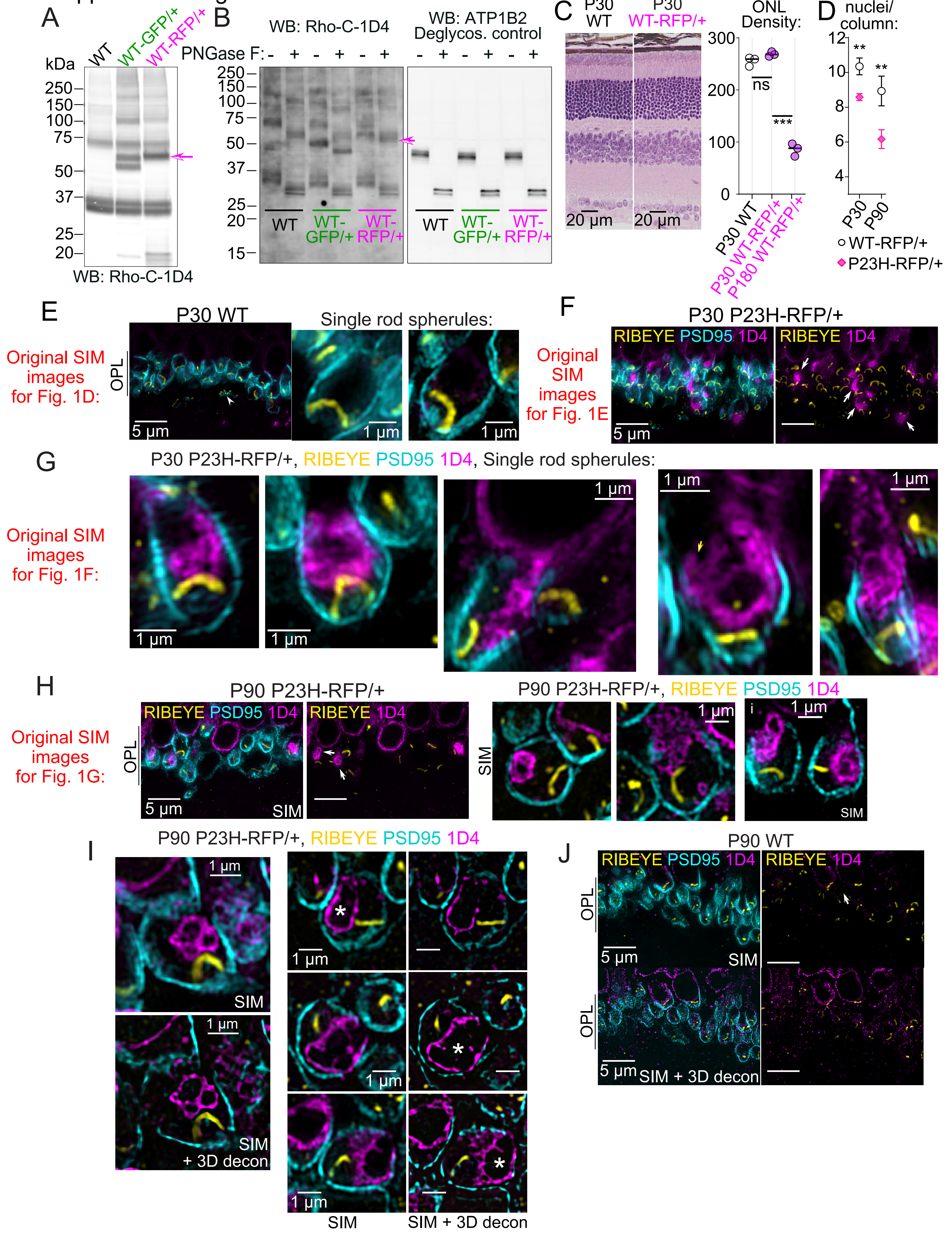

Supplemental Figure 2

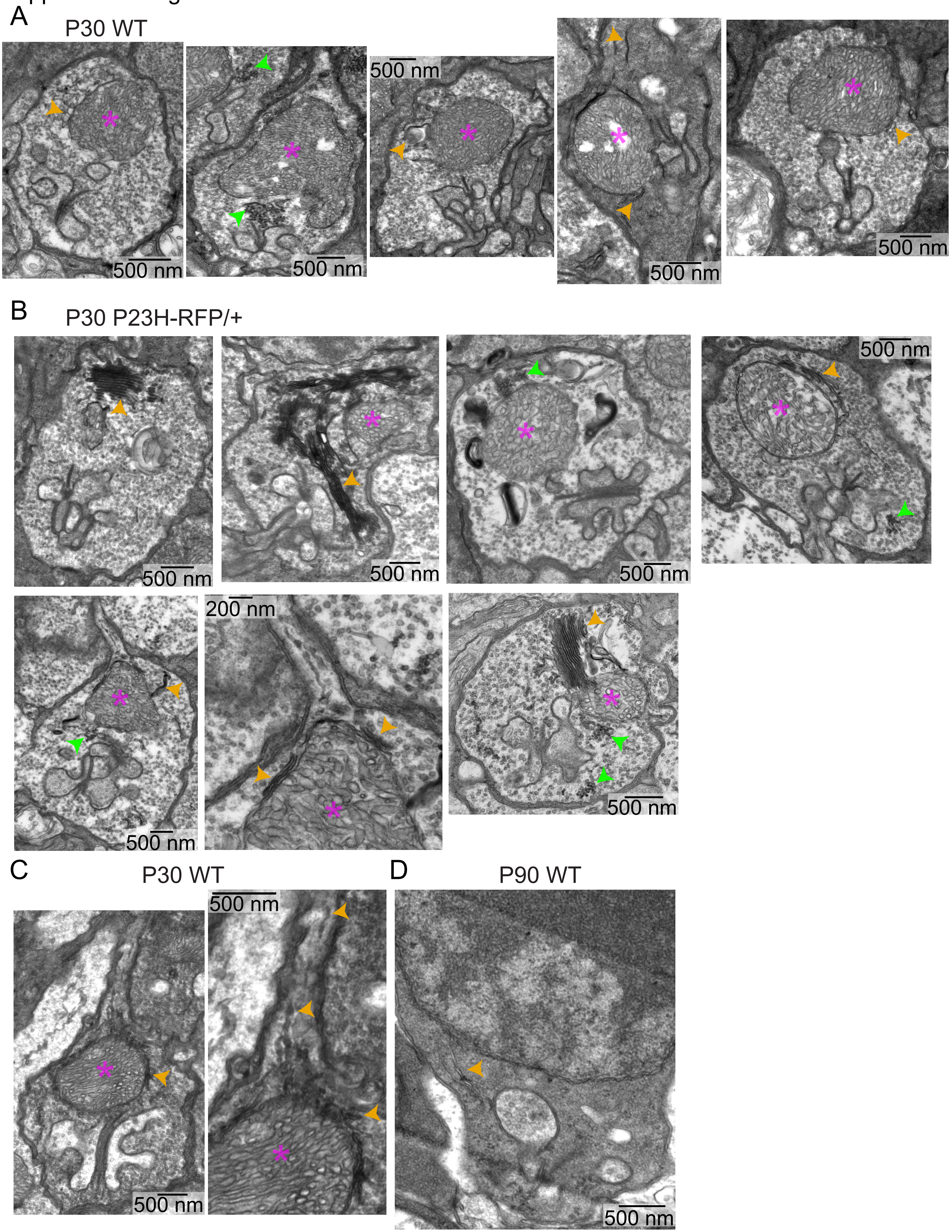

### Supplemental Figure 3

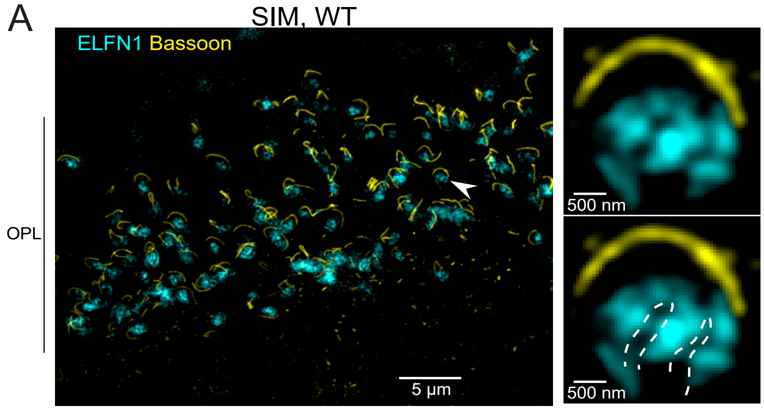


B Normalized TMT-MS values and P-values corresponding to Fig. 4 C and E and other synaptic proteins:

| P30 | Relative Protein Abundance, normalized | | | | | | | | |
|---|---|---|---|---|---|---|---|---|---|
| | WT 1 | WT 2 | WT 3 | P23H-RFP/+ 1 | P23H-RFP/+ 2 | P23H-RFP/+ 3 | P-Value | Adjusted P-Value | # of unique peptides |
| Rho | 1.00439 | 1.1238 | 0.8718 | -1.2161 | -0.904 | -1.2694 | 1E-06 | 0.000374 | 5 |
| Prph2 | 1.07941 | 0.98647 | 0.9341 | 0.3385 | 0.48175 | 0.09223 | 0.0004 | 0.015837 | 12 |
| Pdc | 1.0128 | 0.95589 | 1.03132 | 1.14305 | 1.11667 | 0.98622 | 0.1606 | 0.387183 | 15 |
| Reep6 | 1.0331 | 0.99141 | 0.97549 | 1.33455 | 1.26951 | 1.08723 | 0.0134 | 0.104377 | 7 |
| Tulp1 | 0.99378 | 1.0477 | 0.95853 | 0.79696 | 0.67673 | 0.63401 | 0.0012 | 0.026752 | 19 |
| Elfn1 | 0.94637 | 1.1101 | 0.94354 | 0.59435 | 0.66627 | 0.40601 | 0.0014 | 0.029664 | 5 |
| Grm6 | 0.93813 | 1.10785 | 0.95402 | 0.99374 | 0.58454 | 0.83809 | 0.123 | 0.337101 | 6 |
| Dag1 | 1.02519 | 0.9734 | 1.00141 | 0.92036 | 1.10252 | 1.15252 | 0.3983 | 0.626117 | 6 |
| Dmd | 0.98849 | 1.00599 | 1.00551 | 0.80889 | 0.90406 | 0.81128 | 0.0055 | 0.062718 | 24 |
| Egflam | 1.01361 | 1.00522 | 0.98118 | 0.94404 | 0.7147 | 0.81928 | 0.0277 | 0.153483 | 6 |
| Erc1 | 0.99789 | 1.00214 | 0.99997 | 1.00357 | 1.00524 | 1.00283 | 0.07 | 0.247041 | 10 |
| Rims2 | 0.94394 | 1.01866 | 1.0374 | 0.77752 | 0.89286 | 0.73811 | 0.0086 | 0.080247 | 11 |
| Bsn | 0.98681 | 1.01382 | 0.99939 | 0.92102 | 0.94409 | 0.82331 | 0.045 | 0.196877 | 46 |
| Guca1b | 1.00207 | 0.99645 | 1.00148 | 0.96703 | 0.92383 | 0.95247 | 0.0032 | 0.046721 | 5 |
| Cabp4 | 0.99105 | 1.00866 | 1.00028 | 0.99855 | 0.97512 | 0.9965 | 0.2309 | 0.470574 | 2 |
| Dlg4 | 0.98434 | 1.00392 | 1.01176 | 0.88817 | 0.86946 | 0.82855 | 0.0048 | 0.057393 | 19 |
| Gpr179 | 1.0017 | 0.99808 | 1.00022 | 0.99073 | 0.99447 | 0.98772 | 0.0065 | 0.069647 | 18 |
| Cacna1f | not detected | | | | | | | | |
| Cacna2d4 | 1.00027 | 0.99941 | 1.00032 | 1.00152 | 1.00202 | 0.99604 | 0.9494 | 0.980638 | 5 |
| Dnm1 | 0.99935 | 1.00033 | 1.00032 | 1.00046 | 0.99888 | 0.997 | 0.4063 | 0.633663 | 28 |
| Dnm3 | 0.99936 | 1.00022 | 1.00042 | 1.00031 | 0.99766 | 1.00075 | 0.7734 | 0.889683 | 27 |
| Cplx3 | 0.99824 | 1.00346 | 0.9983 | 0.99575 | 0.99855 | 0.99026 | 0.1001 | 0.299862 | 10 |
| Syt1 | 0.99984 | 1.00013 | 1.00003 | 0.9988 | 0.99947 | 0.99747 | 0.2604 | 0.499433 | 13 |
| Syt7 | 1.00039 | 0.99949 | 1.00013 | 1.00523 | 0.99974 | 0.99968 | 0.4917 | 0.69975 | 4 |

| P90 | Relative Protein Abundance, normalized | | | | | | | | | |
|---|---|---|---|---|---|---|---|---|---|---|
| | WT 1 | WT 2 | WT 3 | P23H-RFP/+ 1 | P23H-RFP/+ 2 | P23H-RFP/+ 3 | P23H-RFP/+ 4 | P-Value | Adjusted P-Value | # of unique peptides |
| Rho | 0.92643 | 1.01164 | 1.06192 | -0.3345 | -0.4548 | -0.2541 | -0.6513 | 8E-07 | 0.00026 | 6 |
| Prph2 | 0.96375 | 0.96499 | 1.07126 | 0.50809 | 0.38464 | 0.47112 | 0.2157 | 7E-05 | 0.00494 | 11 |
| Pdc | 0.96611 | 1.04981 | 0.98409 | 1.48983 | 1.31166 | 1.45948 | 1.31671 | 0.0002 | 0.00865 | 16 |
| Reep6 | 0.97508 | 0.97794 | 1.04698 | 1.31938 | 1.33736 | 1.26472 | 1.34375 | 0.0001 | 0.006 | 7 |
| Tulp1 | 0.95793 | 0.97732 | 1.06473 | 0.64773 | 0.7599 | 0.78762 | 0.84845 | 0.0041 | 0.04401 | 21 |
| Elfn1 | 0.97805 | 1.02672 | 0.99522 | 0.77199 | 0.65646 | 0.84868 | 0.84301 | 0.0054 | 0.05191 | 5 |
| Grm6 | 1.14729 | 0.76821 | 1.0845 | 0.51704 | 0.48131 | 0.43289 | 0.69262 | 0.0023 | 0.03353 | 9 |
| Dag1 | 1.06117 | 1.00332 | 0.93551 | 1.13528 | 1.23299 | 1.22109 | 1.29115 | 0.0038 | 0.04268 | 7 |
| Dmd | 0.99372 | 0.96853 | 1.03777 | 0.91641 | 1.02678 | 0.82786 | 1.04276 | 0.4749 | 0.66894 | 19 |
| Egflam | 0.82856 | 1.13925 | 1.03221 | 0.83232 | 0.56662 | 0.76776 | 0.57212 | 0.0132 | 0.08531 | 6 |
| Erc1 | 0.99966 | 0.99875 | 1.0016 | 1.00991 | 1.01131 | 1.01101 | 1.01892 | 0.0022 | 0.03234 | 7 |
| Rims2 | 0.96643 | 1.04059 | 0.99299 | 0.69649 | 0.60292 | 0.77521 | 0.54496 | 0.0007 | 0.01592 | 4 |
| Bsn | 0.91717 | 0.98127 | 1.10156 | 0.9271 | 1.04839 | 0.98837 | 1.24127 | 0.5574 | 0.73452 | 52 |
| Guca1b | 1.00359 | 0.99894 | 0.99747 | 0.97673 | 0.95402 | 0.96205 | 0.96541 | 0.0002 | 0.00717 | 3 |
| Cabp4 | 0.99924 | 1.00474 | 0.99601 | 0.98042 | 0.97498 | 0.9792 | 0.98287 | 0.0002 | 0.00758 | 3 |
| Dlg4 | 0.98008 | 0.99457 | 1.02537 | 0.88184 | 0.85831 | 0.78408 | 0.90343 | 0.0132 | 0.08531 | 17 |
| Gpr179 | 0.99911 | 0.99741 | 1.00347 | 0.99154 | 0.99493 | 0.99092 | 0.99854 | 0.0616 | 0.20263 | 15 |
| Cacna1f | 13.5825 | 13.5835 | 13.3883 | 13.1042 | 13.3277 | 13.4593 | 13.2791 | 0.0406 | 0.15959 | 0 |
| Cacna2d4 | 19.0163 | 19.0197 | 19.0496 | 19.0631 | 18.9873 | 19.0146 | 18.9102 | 0.4864 | 0.67556 | 7 |
| Dnm1 | 23.8637 | 23.8824 | 23.9203 | 23.8982 | 23.9636 | 23.9532 | 23.8513 | 0.554 | 0.73173 | 28 |
| Dnm3 | 23.9432 | 23.9473 | 23.9314 | 23.9633 | 23.9359 | 23.9815 | 23.7351 | 0.5931 | 0.76012 | 23 |
| Cplx3 | 24.8706 | 24.8483 | 24.8732 | 24.7183 | 24.8055 | 24.8908 | 25.031 | 0.9722 | 0.98759 | 10 |
| Syt1 | 26.9321 | 26.9107 | 26.8771 | 26.9125 | 26.8963 | 26.9784 | 26.7683 | 0.7615 | 0.87062 | 17 |
| Syt7 | 17.5321 | 17.4769 | 17.5615 | 17.5261 | 17.5463 | 17.5066 | 17.6104 | 0.6096 | 0.77195 | 4 |

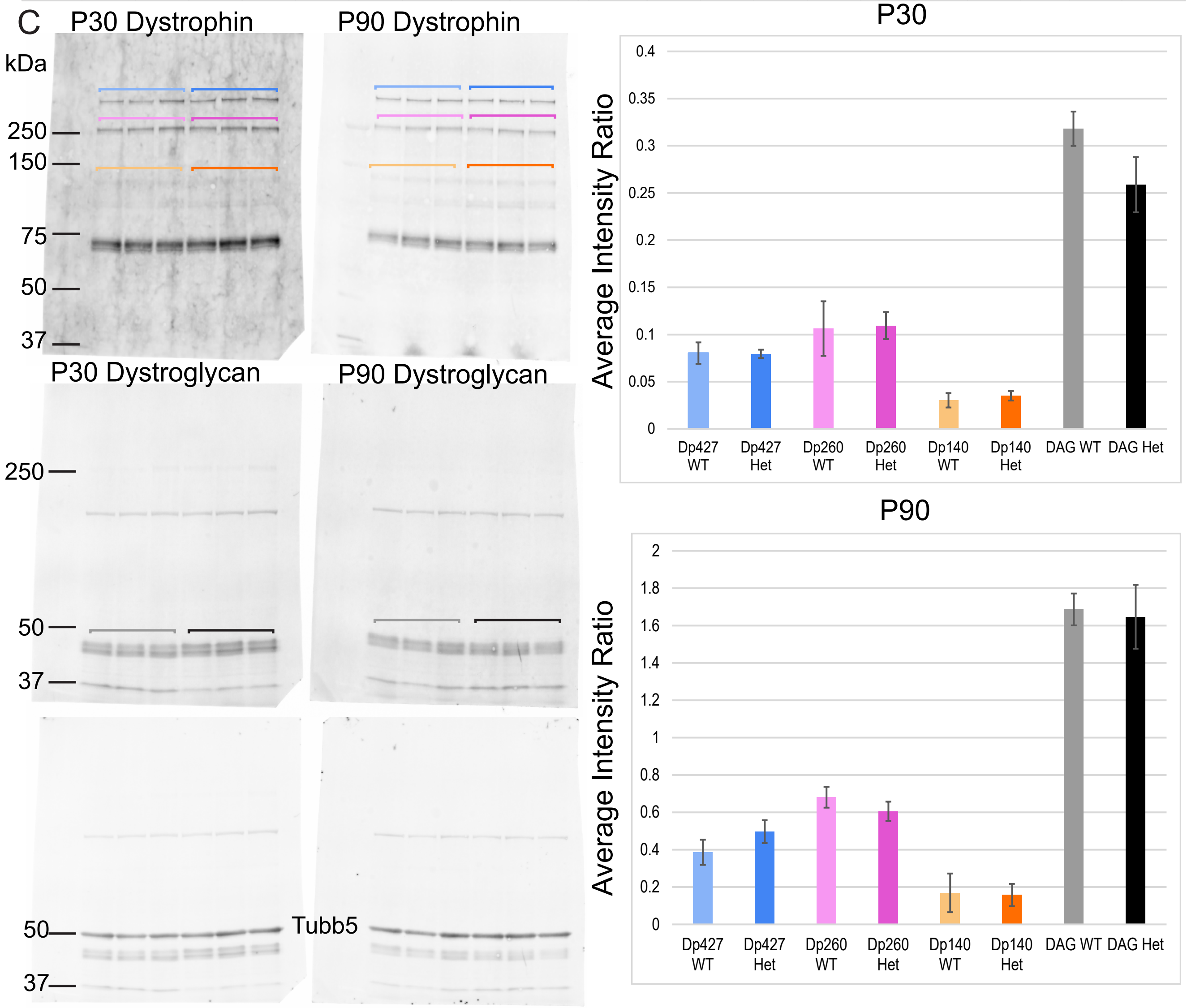

Supplemental Figure 4

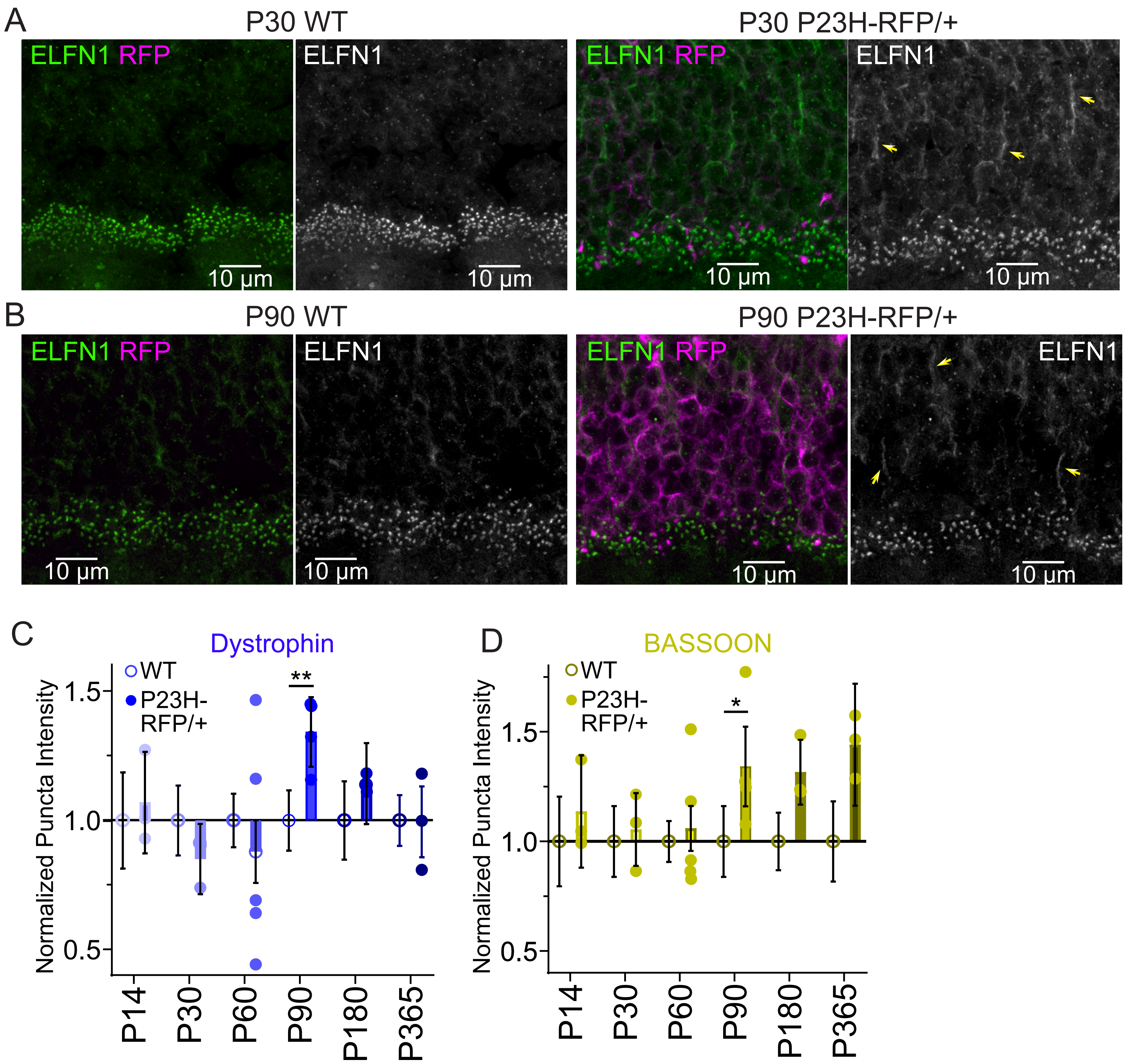

Supplemental Figure 5

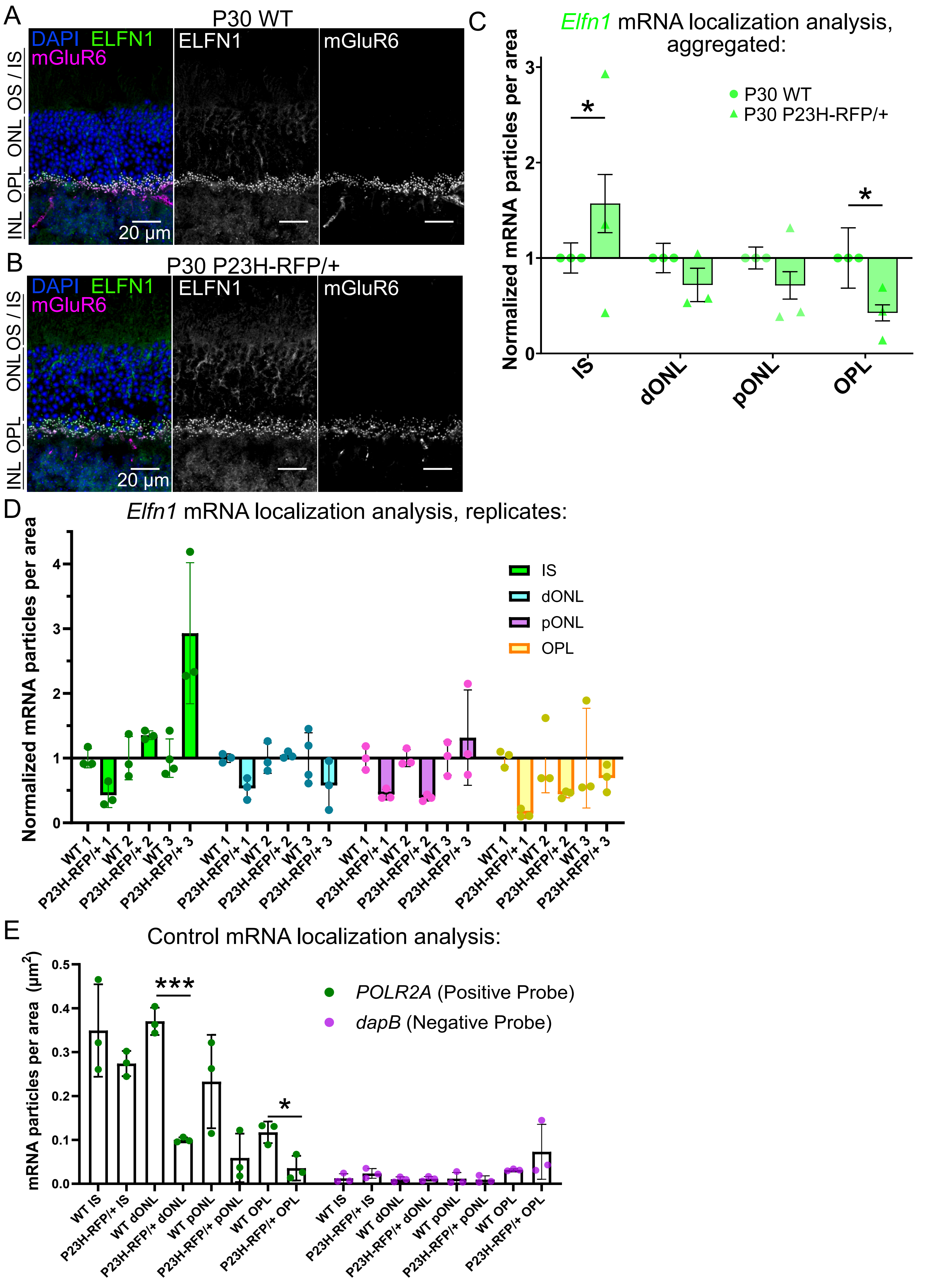

Supplemental Figure 6

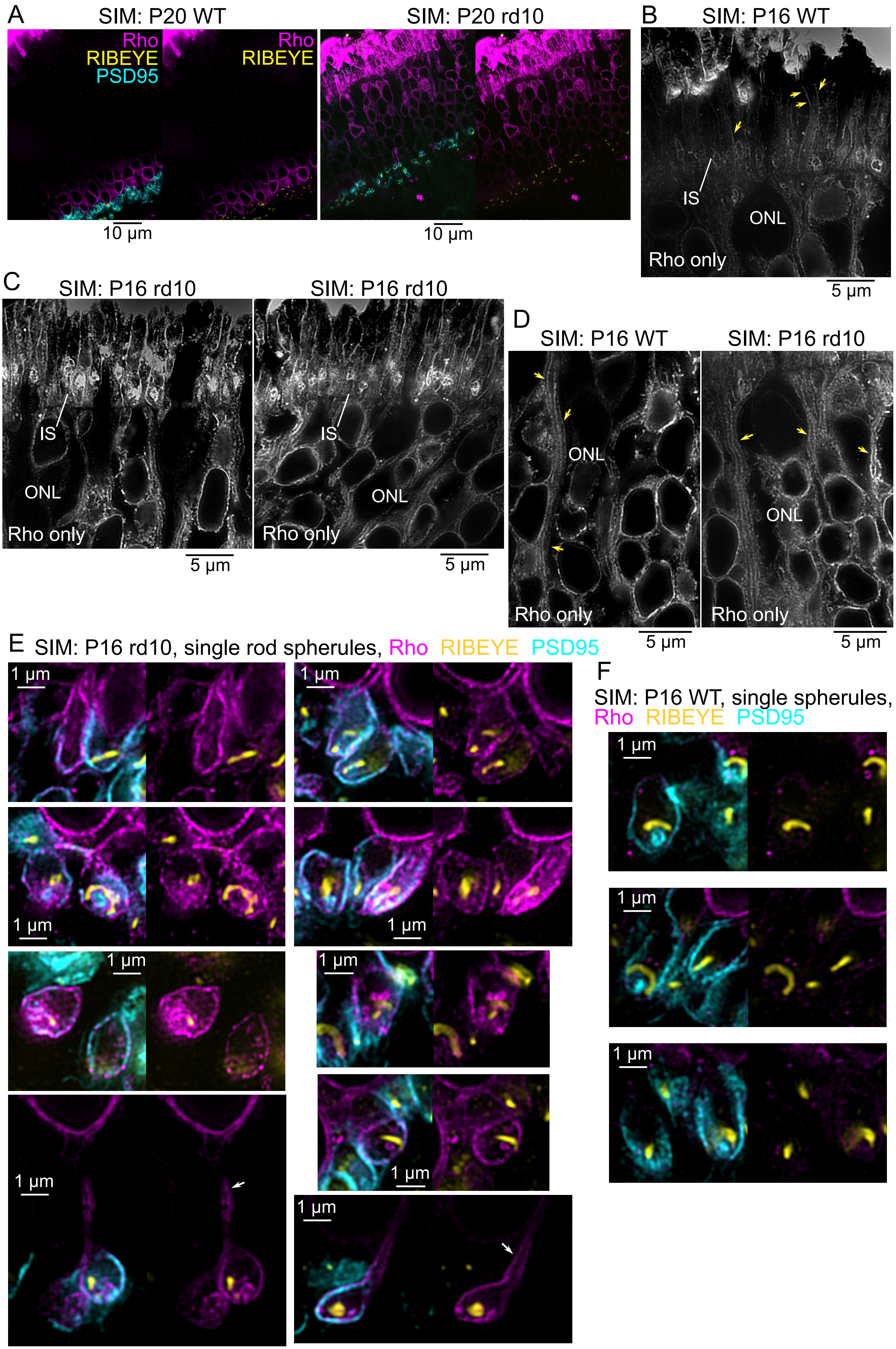
